# Single cell RNA-seq analysis of spinal locomotor circuitry in larval zebrafish

**DOI:** 10.1101/2023.06.06.543939

**Authors:** Jimmy J. Kelly, Hua Wen, Paul Brehm

## Abstract

Identification of the neuronal types that form the specialized circuits controlling distinct behaviors has benefited greatly from the simplicity offered by zebrafish. Electrophysiological studies have shown that additional to connectivity, understanding of circuitry requires identification of functional specializations among individual circuit components, such as those that regulate levels of transmitter release and neuronal excitability. In this study we use single cell RNA sequencing (scRNAseq) to identify the molecular bases for functional distinctions between motoneuron types that are causal to their differential roles in swimming. The primary motoneuron (PMn) in particular, expresses high levels of a unique combination of voltage-dependent ion channel types and synaptic proteins termed functional ‘cassettes’. The ion channel types are specialized for promoting high frequency firing of action potentials and augmented transmitter release at the neuromuscular junction, both contributing to greater power generation. Our transcriptional profiling of spinal neurons further assigns expression of this cassette to specific interneuron types also involved in the central circuitry controlling high speed swimming and escape behaviors. Our analysis highlights the utility of scRNAseq in functional characterization of neuronal circuitry, in addition to providing a gene expression resource for studying cell type diversity.

## Introduction

Functional and anatomical studies of spinal circuitry among the vertebrates have formed the basis of our understanding of control over stereotypic movements (Goulding 2009, Grillner & Jessell 2009). Investigation into movement control continues to benefit from larval zebrafish, which represent a greatly simplified system for resolving the underlying spinal circuitry (Fetcho & Liu 1998, Drapeau et al 2002, Lewis & Eisen 2003, Fetcho et al 2008). Study of circuitry in zebrafish also provides the unique opportunity to trace locomotory circuitry from sensory initiation to the final motor output (Fetcho 1991, Koyama et al 2011, Fidelin & Wyart 2014, Berg et al 2018). Fortuitously, despite the evolutionary distance between fish and mammals, many classes of spinal interneurons involved in movement control are conserved between species, heightening the potential significance of zebrafish circuitry analysis (Grillner 2003, Goulding 2009).

As a new approach towards circuitry analysis, we turned to scRNAseq. For this purpose, we developed a method for isolation and dissociation of spinal cords from 4 days post fertilization (dpf) zebrafish, an age at which much of the anatomy and physiology of swim control has been published (Bhatt et al 2007, McLean et al 2007, Fetcho et al 2008, Liao & Fetcho 2008, McLean et al 2008, Satou et al 2009, Menelaou & McLean 2012, Wang & Brehm 2017, Bello-Rojas et al 2019, Menelaou & McLean 2019, Kishore et al 2020, Satou et al 2020, Wen et al 2020). Our analysis has provided identification of major classes of neuronal and glial types. Of those neuronal types, many are known to be directly involved in locomotory circuitry either in mouse, zebrafish, or both. Using the markers revealed by the transcriptome analysis, we validated a number of key interneuron types, previously shown to be present in zebrafish locomotion circuit. In addition, we identified a new excitatory interneuron type that has a unique transmitter phenotype among interneurons.

Our interest in applying scRNAseq methodology to spinal neurons of zebrafish went beyond identifying transcriptional markers. Rather, we sought to use the neuron-specific transcriptomics to mine for distinctions in the ion channels and synaptic players that are associated with specialized circuit function. Many physiological studies have indicated fundamental differences in excitability and synaptic transmission among neurons controlling escape behavior versus those controlling slow to moderate rhythmic swimming speeds (Bhatt et al 2007, Liao & Fetcho 2008, McLean et al 2008, Satou et al 2009, Menelaou & McLean 2012, Menelaou & McLean 2019, Kishore et al 2020, Satou et al 2020). The signaling molecules causal to these differences among spinal neuron types are largely unknown. To address this outstanding question, we compared different types of motor neurons, where two well studied subtypes serve different roles. The PMns control the strongest contraction that provide for the single powerful bend, initiating escape, and participate in rhythmic swimming at only the highest speeds. By contrast, the SMns collectively regulate the range of slower rhythmic swimming (McLean et al 2007, Gabriel et al 2011, Wang & Brehm 2017). Paired patch clamp recordings, possible only in zebrafish, have characterized these Mn types as functional bookends. The PMns fire action potentials at ultrahigh frequency and the neuromuscular synaptic responses are able to follow with fidelity, whereas the SMns respond with lower frequency action potential firing and synaptic transmission at the neuromuscular junction and are subject to frequent failures (Wang & Brehm 2017, Wen et al 2020).

Our scRNAseq analysis, using Mn subtype specific markers that we validate in this paper, has provided candidates for serving these functional differences. First, the PMns express a trio of unique voltage-dependent ion channels, seen only at very low levels in the SMns, that are tailored for high frequency transmission. The same fast ion channel cassette also enriched in two well-characterized interneuron types that control firing of the PMns and are directly involved in the fast swimming and escape behavior. Second, the PMns also express significantly higher transcript levels of several key proteins involved in exocytosis, collectively termed a synaptic cassette. Thus, scRNAseq offers a new means to interrogate spinal circuitry through assignment of specialized signaling molecules. This application may also prove useful to understanding specialized circuits within the central nervous system (CNS) of mammals.

## Results

### Transcriptional profiling of the larval zebrafish spinal cord

scRNAseq was performed on 4 dpf larval zebrafish, an age corresponding to many studies of spinal circuity and electrophysiological analysis of neuronal control over swimming behavior (Bhatt et al 2007, Liao & Fetcho 2008, McLean et al 2008, Satou et al 2009, Menelaou & McLean 2012, Wang & Brehm 2017, Menelaou & McLean 2019, Kishore et al 2020, Satou et al 2020, Wen et al 2020). For each of two duplicate experiments, approximately 150 spinal cords were isolated, followed by dissociation into single cells. In each case, ~10,000 to 15,000 spinal cord cells were sequenced to a depth of 40,000 reads per cell and aligned to an improved zebrafish reference genome that is more inclusive of 5’ and 3’ untranslated regions (Lawson et al 2020);see Methods). After applying standard quality control filters, data integration resulted in a total of 11762 spinal cord cells from the combined datasets.

Graph-based, unsupervised clustering of the entire spinal cord transcriptome gave rise to 30 clusters, 27 of which were readily identifiable on the basis of established neuronal or glial markers forming the two broad categories of cell types (Fig. 1A). The neuronal markers were *elavl4* and *snap25a* (Fig. 1B) and glia markers were *gfap, slc1a2b, myrf and sox10* (Fig. 1C). The glial cells (46% of total cells) fell into two non-overlapping groups, corresponding to astrocytes/radial glia (*gfap+/slc1a2b+),* and oligodendrocytes (*sox10+/myrf*+) (Fig. 1C). Both glial types were composed of multiple clusters, indicating further diversity. The remaining 3 clusters (clusters 6, 11, and 17, corresponding to 11% of total cells) showed mixed expression of neuronal and glial markers. It’s unclear whether this population reflects a true cell type or instead a group of unremoved doublets. Given that these clusters could not be assigned to either neurons or glia with confidence they were excluded from the subsequent analysis. Additionally, while the glia data are available as a resource, they were not analyzed further in this study.

**Figure 1.**
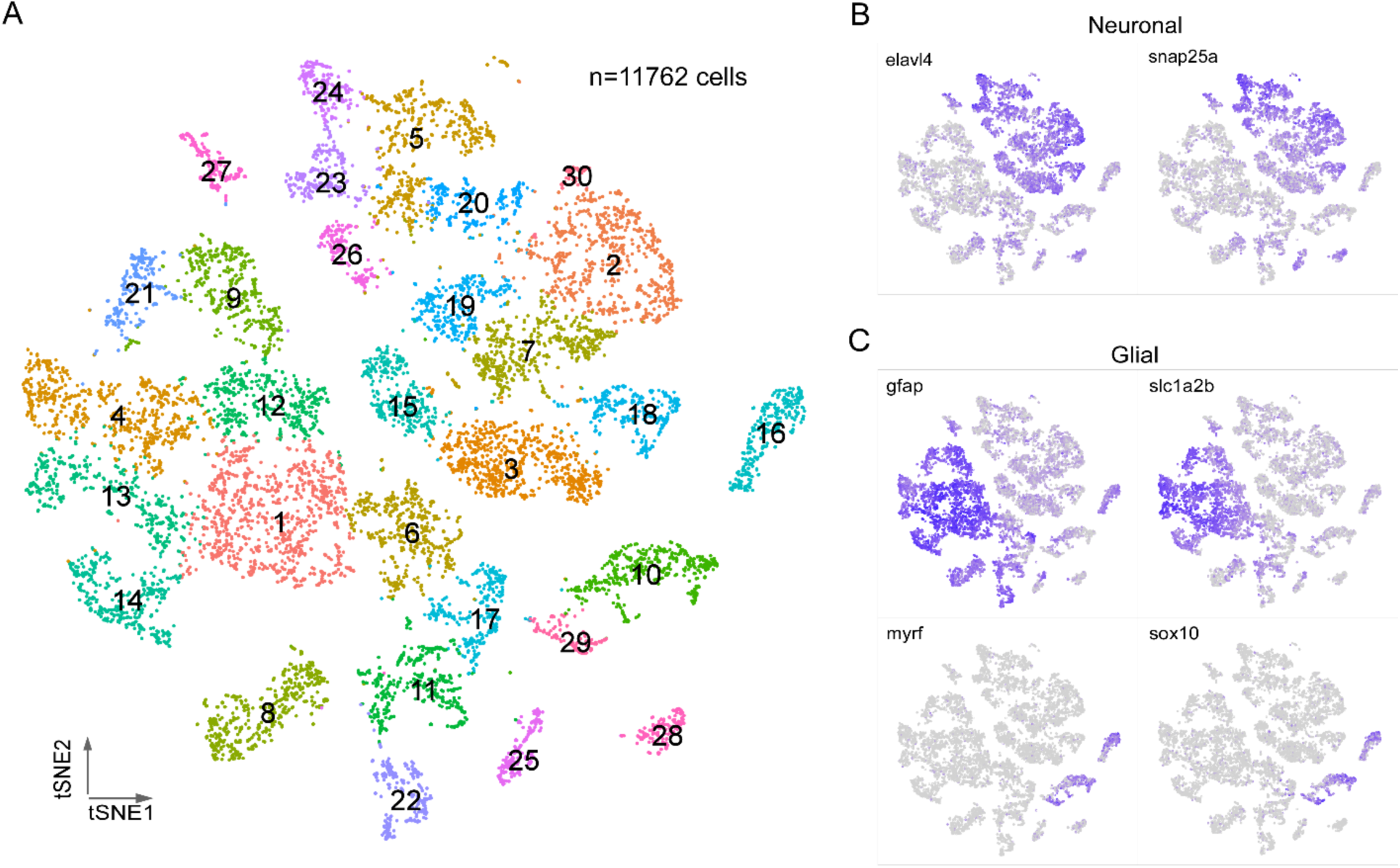
Transcriptional profiling of larval spinal cord. A. Visualization of 4 dpf spinal cord cells using t-distributed stochastic neighbor embedding (t-SNE). Each dot is a cell and each arbitrary color corresponds to a single cluster. The clusters are individually numbered and the total number of cells indicated. B-C. Feature plots for 2 neuron markers (B) and 4 glial makers (C). Two sets markers are shown to distinguish the two broad types of glial cells, *gfap* and *slc1a2b* for astrocytes/radial glia (C, top), *myrf* and *sox10* for oligodendrocytes (C, bottom).

### Transcriptional profiling of spinal cord neurons

The neuronal population identified by *elavl4+*/*snap25a*+ expression was re-grouped into 33 clusters using Seurat (Fig. 2A). To assign individual neuronal identities to these transcriptome clusters, we used a combination code that relied on the co-expression of neurotransmitter biosynthesis/transporter genes along with differentially expressed marker genes (DEGs; Fig. 2; Table 1; Supplementary Table 1). The first code provided clear separation of neuronal populations into four principal categories; glutamatergic (*slc17a6a+/slc17a6b+*), glycinergic (*slc6a5+*), GABAergic (*gad1b+/gad2+*) or cholinergic (*chata+/slc18a3a+*) types (Fig 2B, Table 1). The second code relied on not only established markers for both zebrafish and mouse spinal neurons, but also new markers identified in this study. Generating the list of candidate markers was aided by previous studies that profiled the neurotransmitter identity of morphologically distinct neurons in the larval spinal cord (Bernhardt et al 1990, Hale et al 2001, Higashijima et al 2004a, Higashijima et al 2004c). In addition, since many aspects of the transcriptional program that establish the spinal neuronal circuit have been shown to be evolutionarily conserved among vertebrates (Kiehn & Kullander 2004, Goulding 2009, Grillner & Jessell 2009), we cross-referenced our data with recent mouse spinal cord sequencing data to search for homologous marker genes (Delile et al 2019, Blum et al 2021).

**Figure 2.**
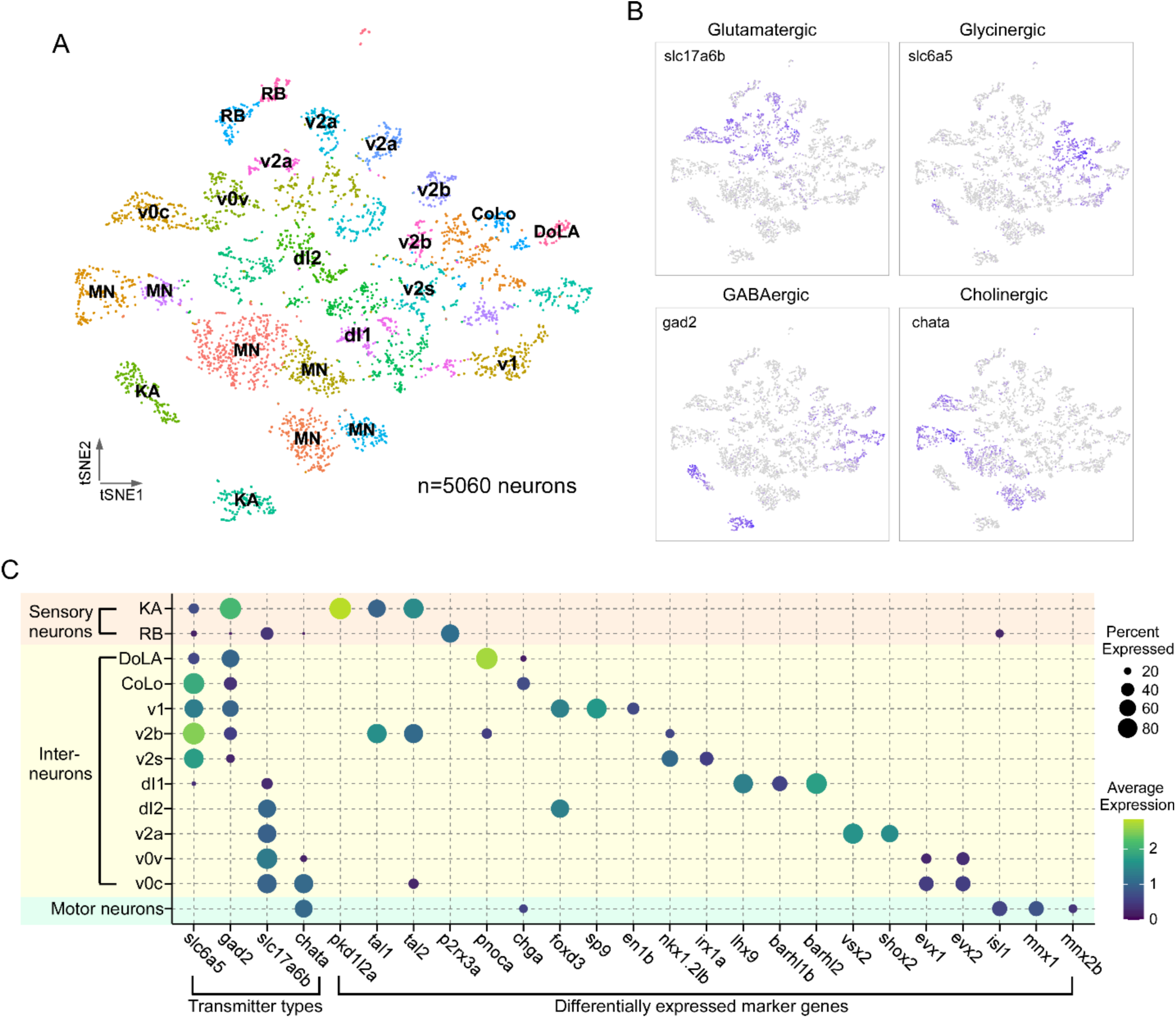
Transcriptional profiling of larval spinal cord neurons. A. Visualization of neuronal populations for 4 dpf spinal cord using t-SNE. Each dot is a cell and each arbitrary color represents a cluster. Cell type identity assigned to each cluster utilized the combination code of neurotransmitter phenotype, marker genes and morphological labeling. B. Feature plots for the four major classes of excitatory and inhibitory neurotransmitter genes. Vesicular glutamate transporter vGlut2 (*slc17a6b*) was used for glutamatergic neurons; glycine transporter glyt2 (*slc6a5*) for glycinergic neurons; glutamate decarboxylase (*gad2*) for GABAergic neurons; choline acetyltransferase *(chata)* for cholinergic neurons. C. Dot plot showing neuronal cell identity versus markers used for assignment. Dot size indicates the percentage of cells in the cluster showing expression of the indicated marker and color scale denotes the average expression level. For visual clarity, dot sizes below 15 percent expressed are omitted.

**Table 1.** Combination codes used for assigning cell types to clusters.

| **Cell Type** | **Neurotransmitter** | **Markers** | **% of neurons** |
| --- | --- | --- | --- |
| **KA** | GABA | pkd1l2a, tal1,tal2 | 7.0 |
| **RB** | glutamate | p2rx3a | 3.8 |
| **DoLA** | GABA | pnoca | 1.4 |
| **CoLo** | glycine | chga | 1.1 |
| **v1** | glycine/GABA | sp9, foxd3, en1b | 4.1 |
| **v2b** | glycine | tal1, tal2 | 3.6 |
| **v2s** | glycine | nkx1.2lb, irx1a | 2.7 |
| **dl1** | glutamate | lhx9, barhl1b, barhl2 | 1.7 |
| **dl2** | glutamate | foxd3 | 3.7 |
| **v2a** | glutamate | vsx2, shox2 | 6.2 |
| **v0v** | glutamate | evx1,evx2 | 4.0 |
| **v0c** | glutamate/acetylcholine | evx1, evx2, | 4.2 |
| **Mns** | acetylcholine | isl1, mnx1, mnx2b | 26.8 |

Using the combination code, we identified specific classes of sensory neurons, Mns and interneurons (Fig. 2C, Table 1). Two types of sensory neurons, the glutamatergic Rohon-Beard (RB) cells and GABAergic Kolmer-Agduhr (KA; also referred to as cerebrospinal fluid-contacting neurons (CSF-cN) cells) are each represented by two distinct clusters (Fig. 2A & C). Cholinergic clusters formed a prominent group corresponding principally to Mns. Clusters of interneurons corresponded to the inhibitory v1, v2b and v2s, and excitatory v0v, v2a and dl1, and dl2 types (Fig. 2A & C). The relative abundance of the cell types associated with individual clusters (Table 1) were consistent with those published for *in vivo* labeling experiments (Higashijima et al 2004b, Kimura et al 2006, Ampatzis et al 2014, Gerber et al 2019, Satou et al 2020), suggesting that our preparation protocol and analysis are robust and unbiased in recovering spinal cell populations. Of the 33 clusters in our neuronal dataset we were able to assign identities to 22 clusters with confidence. There are a number of cell types described for in larval zebrafish, such as the excitatory v3 interneuron or the inhibitory v0d interneuron (Satou et al 2020, Bohm et al 2022) that we were unable to identify, likely due to lower expression levels of canonical markers at this developmental stage. It is probable that these interneuron populations are present in unassigned clusters.

Our analysis also resolved subtypes within neuronal classes. For example, the two clusters were assigned as KA neurons, that shared a common set of markers for the cerebrospinal fluid contacting interneuron *(pkd1l2a/pkd2l1*; Fig. 3A). However, assignment to subtype can be made on the basis of differential expression of *sst1.1* versus *urp1*, previously shown to label KA’ and KA’’ functional groups, respectively (Fig. 3A) (Djenoune et al 2017, Yang et al 2020).

**Figure 3.**
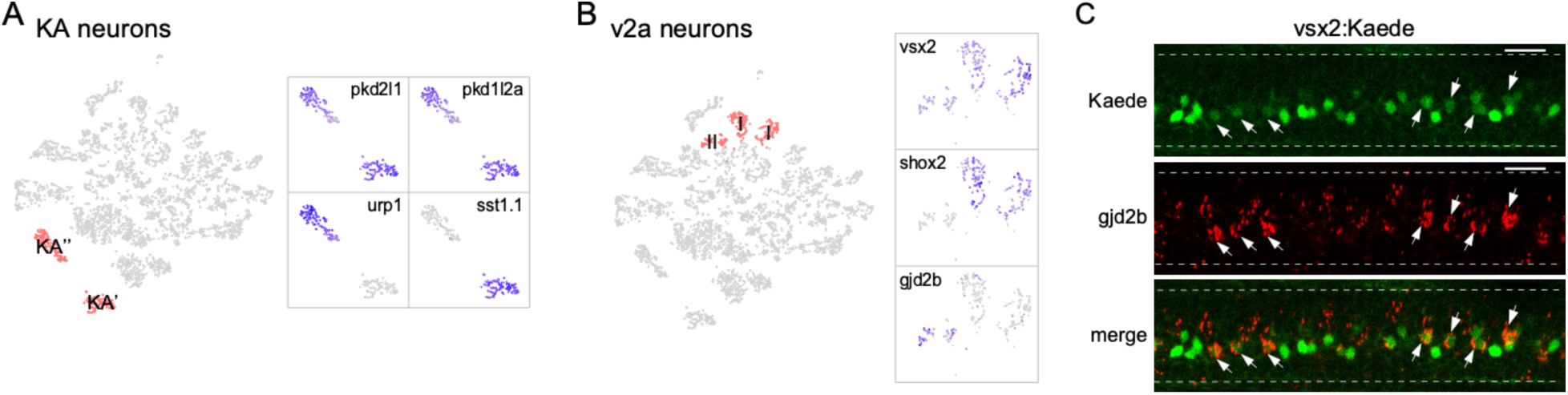
Diversity in neuronal types. A. Zoomed feature plots for *pkd2l1, pkd1l2a, urp1 and sst1.1* that differentiate the KA’ and KA’’ neurons (right). The two clusters correspond to KA’ and KA’’ neurons indicated in the neuronal t-SNE projection (left, in red). B. Zoomed feature plots for *vsx2, shox2 and gjd2b* that differentiate the Type I and Type II v2a neurons (right). The three clusters corresponding to v2a interneurons indicated in the neuronal t-SNE projection (left, in red). C. Representative *in situ* hybridization images showing enriched expression of *gjd2b* in Type II v2a (arrows) in a Tg(vsx2: Kaede) transgenic fish. The two sub-groups of v2as were discerned with different levels green Kaede fluorescence. n = 8 fish. Scale bar 20 μm. Spinal cord boundary indicated with dashed lines.

Similarly, the glutamatergic v2a interneurons consist of multiple clusters that represented distinct subtypes. This class of interneurons has previously been divided into two sub-populations, Types I and II, based on morphology, molecular and functional heterogeneity (Bhatt et al 2007, McLean & Fetcho 2009, Ampatzis et al 2014, Menelaou et al 2014). One molecular feature differentiating the two types is the higher expression level of both the *vsx2* and *shox2* marker genes in Type I compared to Type II v2a (Kimura et al 2006, Menelaou et al 2014, Hayashi et al 2018, Menelaou & McLean 2019). In our dataset both the *vsx2* and *shox2* expression patterns differed among the v2a clusters (Fig. 3B). The two clusters with strong *vsx2* and *shox2* expression likely represent Type I v2a neurons, while the third cluster with weak *vsx2* expression and no *shox2* expression likely represents Type II (Fig. 3B). For this cluster, we further identified *gjd2b*, the gene encoding the gap junction protein connexin 35.1 δ subunit as an additional marker gene (Fig. 3B). This finding is consistent with previous immunohistochemistry in adult fish demonstrating selective expression *of gjd2* in Type II v2a axons (Carlisle & Ribera 2014, Pallucchi et al 2022). Subtype assignment based on expression patterns of these markers was validated using *in situ* hybridization in larval Tg(vsx2: Kaede) fish (Fig. 3C). Expression of the Kaede fluorescent protein driven by *vsx2* promoter labels Type I v2a with strong fluorescence compared to weakly labeled Type II (Fig. 3C). Probes against *gjd2b* colocalized only with those neurons exhibiting weak fluorescence, confirming its specific expression in the Type II v2a subtype (Fig. 3C). The two subtypes of v2a interneurons both make direct connections with the PMns and are recruited during high speed swimming but differ in terms of connection strength and the type II v2a interneuron is recruited more effectively across the range of high swimming speeds (Kimura et al 2006, Bhatt et al 2007, McLean & Fetcho 2009, Menelaou & McLean 2019). The availability of transcriptomes for these neuronal sub-groups provides an opportunity to further mine for differential molecular features responsible for the functional distinction.

In addition to validating established markers for neuronal types and subtypes, our scRNAseq analysis revealed novel marker genes for identifying interneuron transcriptomes, as shown for the Commissural Local (CoLo) and Dorsal Longitudinal Ascending (DoLA) interneurons.

#### CoLo interneurons

CoLo interneurons provide the fast contralateral inhibition necessary for a successful escape response through direct contact with the PMns (Fetcho & Faber 1988, Liao & Fetcho 2008, Satou et al 2009, Kishore et al 2020). CoLo sends axons ventrally that turn to the contralateral side and split into short ascending and descending projections (Higashijima et al 2004a, Higashijima et al 2004c, Liao & Fetcho 2008, Satou et al 2009, Kishore et al 2020, Satou et al 2020) (Fig. 4A1). We found a small glycinergic cluster that expressed high levels of chromogranin A (*chga*) (Fig. 4A2) that was distinct from the *chga*-enriched cholinergic cluster later assigned to the PMn type (Fig. 5D). Labeling using *chga in situ* hybridization probes revealed a distinct large neuron located rostrally in each hemi-segment in addition to several large-sized motor neurons (Fig. 4A3, see also Fig. 5D). The stereotypical location and the one per hemi-segment stoichiometry of these *chga* labeled interneurons is consistent with that of CoLos (Liao & Fetcho 2008, Satou et al 2009, Kishore et al 2020). To validate its identity, we transiently labeled inhibitory interneurons with EGFP under the control of the *dmrt3a* promotor (Kishore et al 2020, Satou et al 2020). Single GFP labeled CoLos were screened based on morphology (Fig. 4A1) and subsequently shown to co-label with *chga* (Fig. 4A3).

**Figure 4.**
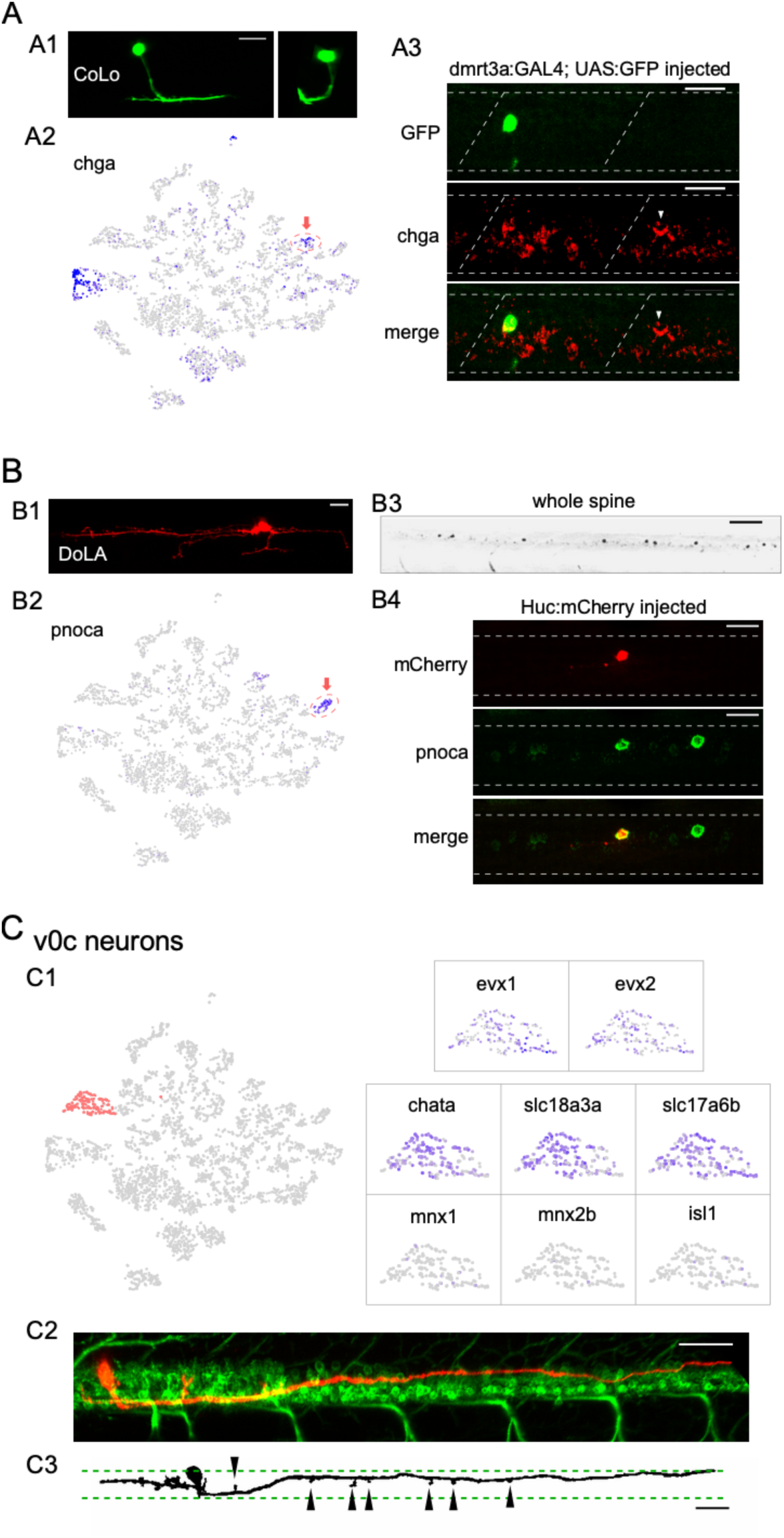
Identification of 3 different interneuron types using the combination code. A. CoLo interneurons. (A1) A CoLo neuron transiently labeled with GFP was identified by its short axons and localized commissural extension. A cross-section provided a clear view of its commissural branching (left). (A2) Feature plot of *chga* in the neuronal t-SNE projection. *chga* expression is localized in the CoLo cluster (red circle with arrow) in addition to a single Mn cluster. (A3) *chga in situ* hybridization probes stained a CoLo labeled with GFP. The CoLo in the neighboring hemi-segment that was not labelled by GFP was also positive (arrowheads). Other positive labeling reflected the PMns (see also Fig. 5D). n= 6 fish. Boundary of spinal cord and segments were indicated (white dash). Scale bar 20 μm. B. DoLA interneurons. (B1) A DoLA transiently labeled with mCherry was identified by its dorsal position and distinct morphology. (B2) Feature plot of *pnoca* in the neuronal t-SNE projection. *pnoca* expression is restricted in the DoLA cluster (red circle with arrow). (B3) *In situ* hybridization of *pnoca* shown for several spinal segments. n= 12 fish. Scale bar 100 μm. (B4) *In situ* hybridization of *pnoca* colocalized with a mCherry-labeled DoLA neuron. n = 7 cells. Scale bar 20 μm. C. v0c interneurons. (C1) Zoomed feature plots for (*evx1, evx2, chata*, *slc18a3a*, *slc17a6b*, *mnx1*, *mnx2b* and *isl1* in the v0c cluster (right). The cluster corresponding to v0c neurons indicated in the neuronal t-SNE projection (left, in red). v0c interneuron cluster is identified by the co-expression of both glutamate (*slc17a6a*) and acetylcholine (*slc18a3a/chata*) pathway genes, and absence of Mn markers (*mnx1/mnx2b/isl1*). (C2) An example of a transiently labelled v0c by mCherry in a 4 dpf Tg(mnx1:GFP) fish. (C3) An example of v0c neurons in gray scale showing the morphology, with boundaries of the motor column (green dash) and enlargements along the axon (arrowheads) indicated. n= 37 fish. Scale bar 50 μm in C2 and C3. Caudal on right and rostral on left.

**Figure 5.**
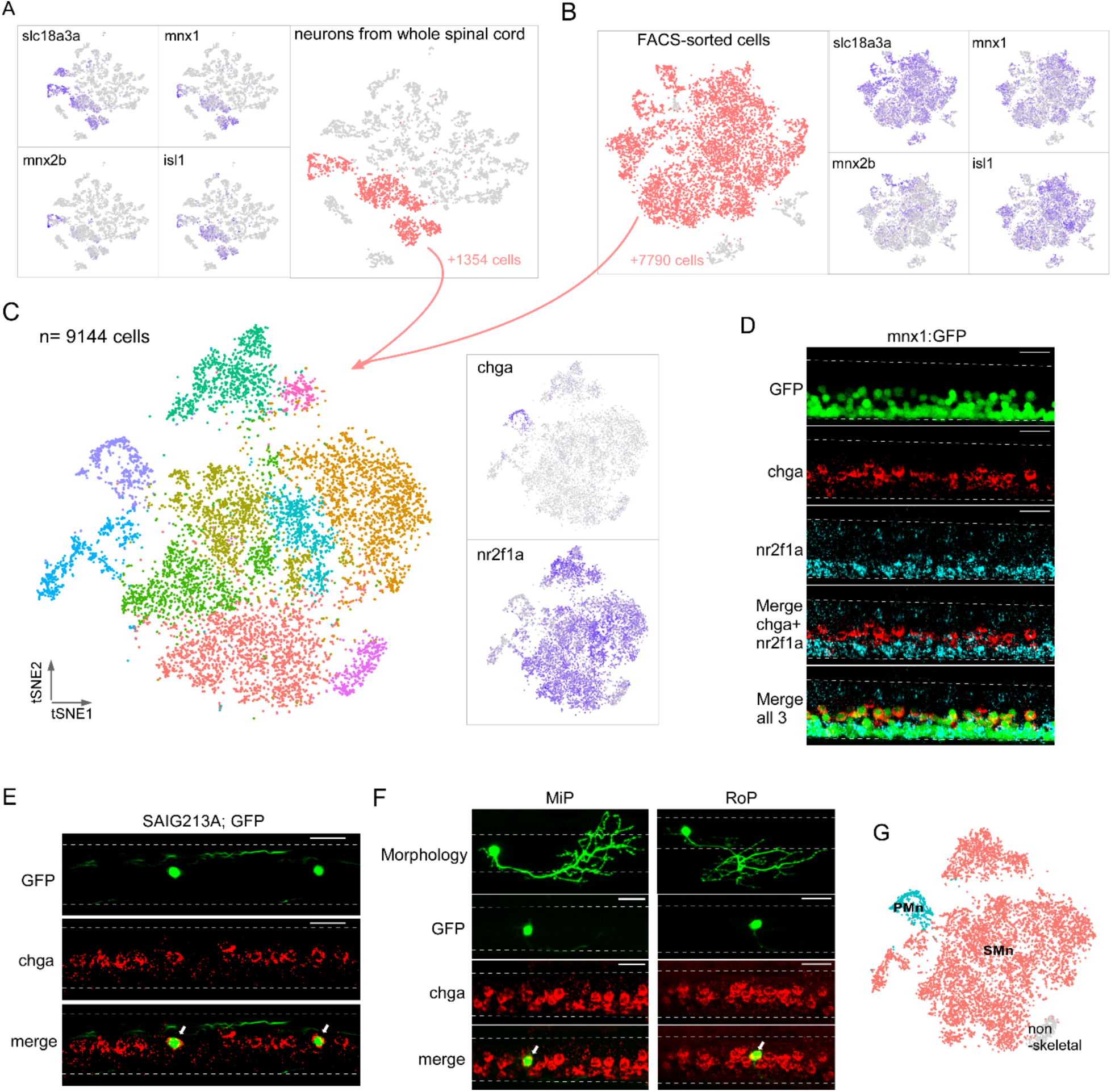
Single cell transcriptional profiling of Mn types at larval stage. Both computational extraction (A) and experimental enrichment (B) approaches were used to isolate Mn populations (red) on the bases of co-expression of acetylcholine transmitter genes (*slc18a3a* shown) and established Mn markers (*mnx1*, *mnx2b* and *isl1*). The total numbers of Mns obtained using each approach indicated. (C) The integrated dataset shown in t-SNE projection, along with feature plots for two marker genes, *chga* and *nr2f1a*. D. Representative *in situ* hybridization images using *chga* and *nr2f1a* probes in a 4 dpf Tg(mnx1:GFP) fish spinal cord. The motor column, indicated by GFP expression, is located ventrally in the spinal cord (top). *chga* and *nr2f1a* signals occupied more dorsal and ventral positions respectively within the motor column (bottom 4 panels). n = 13 fish. E & F. *in situ* hybridization images showing specific expression of *chga* in PMns. Colocalization is shown for GFP-labeled CaP in Tg(SAIG213A;EGFP) fish (indicated by arrows in E, n = 14 fish), and individually labeled MiP and RoP (indicated by arrows in F, n= 4-6 cells). For images in D-F, dorsal is up. Dashed line indicates the spinal cord boundary. Scale bar 20 μm. (G) PMn (cyan), SMn (red) and non-skeletal Mn (gray) assignment.

#### DoLA interneurons

DoLAs are GABAergic inhibitory interneurons located dorsally in spinal cord with well-described morphology (Bernhardt et al 1990, Higashijima et al 2004c) (Fig. 4B1). Our dataset indicated two GABAergic interneuron clusters, one of which corresponded to DoLA neurons with highly specific enrichment of *pnoca* expression (Fig. 4B2). *In situ* hybridization using *pnoca* probes strongly labeled a small number of distinct dorsal neurons located immediately ventral to the domain occupied by the sensory RB cells (Fig. 4B3). Sparse labelling of spinal neurons with a mCherry reporter showed that the neurons positive for *pnoca* projected long ascending axons with short ventral projections, as well as with occasional short descending axons, thus matching the morphological characteristics reported for DoLA interneurons at this stage (Fig. 4B4) (Bernhardt et al 1990, Higashijima et al 2004c, Wells et al 2010). The functional roles played by DoLA neurons have remained elusive. *pnoca* encodes Prepronociceptin, a precursor for several neuropeptides involved in multiple sensory signaling pathways (Martin et al 1998). Its highly specific expression in DoLA suggests that they might function as neuropeptide releasing neurons that modulate sensory functions in larval fish.

Importantly, our analysis also identified a cholinergic/glutamatergic spinal interneuron type not described previously in larval zebrafish. A single cluster in our dataset expressed cholinergic markers that include vesicular acetylcholine transporter vAChT (*slc18a3a*) and choline acetyltransferase (*chata*), but lacks all canonical Mn markers (*mnx1/mnx2b/isl1*) (Fig. 4C1). The cluster also expressed *evx1* and *evx2*, markers associated with interneuron types in the v0 domain (Fig. 2C, Fig. 4C1) (Zagoraiou et al 2009, Juarez-Morales et al 2016). This transcriptional profile suggested that it represented the homologues to the premotor cholinergic v0c interneurons in mouse spinal cord (Zagoraiou et al 2009). Cholinergic interneurons have only recently been shown to be present in adult fish by immunochemistry staining (Bertuzzi & Ampatzis 2018). Similar to their mammalian counterparts, they play roles in modulating Mn excitability (Bertuzzi & Ampatzis 2018). Notably, v0c in larval zebrafish differs from the mouse counterpart on the basis of co-expression of cholinergic and glutamatergic transmitter genes (e.g., *slc17a6b*, Fig. 4C1).

We capitalized on the unique cholinergic phenotype of v0c among interneurons to provide *in vivo* labelling in the spinal cord. For this purpose, we injected a tdTomato reporter driven by the vAChT promoter to sparsely label cholinergic neurons in the 4 dpf spinal cord of Tg(mnx1: GFP) fish (Fig. 4C2). In addition to the Mns, we observed mosaic labeling of an interneuron type with distinct position and morphology. The soma was located near the dorsal boundary of the motor column (Fig. 4C2 & 3) and the axonal processes were either bifurcating (20 out of 37 cells) or purely descending (14 out of 37) or ascending (3 out of 37). The descending processes would reach lengths corresponding to multiple segments within the motor column (Fig. 4C2 & C3, average length > 6 segments). There was an overall lack of secondary branches, but enlargements reminiscent of synaptic boutons in close vicinity of Mn soma were observed along the neurites (Fig. 4C3). Multiple neurons of this type were observed in the same segment even with the sparse labeling approach, suggesting that there are likely to be numerous v0cs in each segment. The morphology and anatomic arrangement are consistent with a role in modulating Mn properties, as has been proposed for v0c in both adult zebrafish (Bertuzzi & Ampatzis 2018) and the mammalian homolog (Zagoraiou et al 2009).

### Subclustering the Mns based on single-cell transcriptomes

We next focused on transcriptome comparison within Mn populations to examine their molecular heterogeneity. As a first approach, we isolated Mn clusters from the whole spinal cord dataset based on the overlap of two sets of marker genes (Fig. 5A). The first set was based on components of the cholinergic pathway that included *slc18a3a,* (Fig. 5A) and *chata* (Fig. 2B). The second set of Mn markers included the transcription factors *mnx1, mnx2b and isl1* (Fig. 2C, Fig. 5A) (Appel et al 1995, Hutchinson & Eisen 2006, Zelenchuk & Bruses 2011, Asakawa et al 2012, Seredick et al 2012). Overall, ~27% of the profiled neuronal single cell transcriptomes corresponded to Mns (1354 cells) (Fig. 5A). As a complementary approach, we also generated samples enriched for a larger number of Mns to increase the statistical power of the clustering analysis. This was achieved through fluorescence activated cell sorting (FACS) of spinal cells prepared from the fluorescent transgenic fish line, Tg(*mnx1:GFP*), that broadly labels Mns (Flanagan-Steet et al 2005, Bello-Rojas et al 2019). Clustering analysis of the sorted data showed that >92% of the FACS sorted cells represent Mns based on the same canonical markers used for whole spinal cord, giving rise to 7790 cells for sub-clustering (Fig. 5B). We integrated the datasets from both the computationally isolated and experimentally purified Mn populations and performed clustering analysis using Seurat. A total of 9144 cells were grouped into 10 clusters (Fig. 5C). A small cluster (3.6% of total) expressing genes *gfra1a* and *tbx3b* (Supplemental Fig. 1) was almost entirely sourced from the FACS sorted datasets (>94%). In addition, it shared markers with a population of non-skeletal muscle Mns recently described in mouse spinal cord scRNAseq datasets (Blum et al 2021). Therefore, this cluster was not included in further comparisons among Mns that control skeletal muscle contraction.

The transcriptionally distinct clusters were next linked to previously known Mn types. Two broad types of Mns have been described for larval zebrafish, the PMn and the SMn, that are commonly distinguished by birth date, progenitor lineage and morphological features such as size, location, and periphery innervation pattern (Eisen et al 1986, Myers et al 1986, Menelaou & McLean 2012, Bello-Rojas et al 2019). There are four large-sized PMns (CaP, MiP, vRoP and dRoP) in each hemi-segment of the spinal cord, each innervating approximately one-quarter of axial muscle target field (Bello-Rojas et al 2019, Wen et al 2020). By contrast, 50-70 SMns are in a more ventral location, and display a gradient of sizes and functional properties (Myers et al 1986, Westerfield et al 1986, McLean et al 2007, Asakawa et al 2012, Wang & Brehm 2017, Wen et al 2020). We examined the top DEGs among the clusters (supplemental Table 2), and found two markers, *chga* and *nr2f1a*, with non-overlapping expression pattern that could reflect this broad classification (Fig. 5C). *chga* was enriched in one distinct cluster, while *nr2f1a* was present in the majority of the remaining cells (Fig. 5C). Taken together, these two mutually exclusive markers represented ~95% of Mn population.

*In situ* hybridization labeling with probes against *chga* and *nr2f1a* revealed spatially segregated Mn groups in the Tg(mnx1:GFP) fish. Specifically, *chga* probes labeled dorsal Mns that were large in size, consistent with them being the primary group (Fig. 5D). *nr2f1a* labeling was absent in these cells, but was distributed in larger number of smaller Mns that were more ventrally located in the motor column (Fig. 5D). These segregated patterns of expression strongly suggested that *chga+* cluster represented the PMns, while *nr2f1a* marked the major SMn populations.

For further validation, we used the *chga* probes for *in situ* hybridization analysis in Tg(SAIG213A; EGFP) fish, in which a single PMn in each hemi-segment, the dorsal projecting CaP, was fluorescently-labeled among all the Mns (Muto & Kawakami 2011, Wen et al 2020). Strong *chga* labeling colocalized with the GFP labeled-CaP in each hemi-segment (Fig. 5E). We also labeled the MiP and RoP Mns in the spinal cord using the sparse labeling approach, and observed high level of *chga* signal in both PMn types using *in situ* hybridization (Fig. 5F). These results firmly established *chga* as the marker for PMns, leaving the *chga* negative clusters representing SMns (Fig. 5G). The number of cells associated with SMn clusters was ~18 fold in excess that of the PMn cluster, consistent with previous cell counts of SMns versus PMns in larval zebrafish (Eisen et al 1986, Myers et al 1986, McLean et al 2007, Menelaou & McLean 2012, Bello-Rojas et al 2019).

### Diversity among SMns

In contrast to the single cluster associated with the PMn type, SMns were composed of multiple transcriptionally distinct groups. Approximately 95% of SMns fall into three groups that were distinguished by differential expression of three marker genes, *foxb1b*, *alcamb* and *bmp16* (Fig. 6A). *In situ* hybridization using the three probes labeled SMns revealed two closely stratified layers that showed differences in dorsal-ventral positioning within the motor column (Fig. 6A). The *foxb1b*+ SMns occupied the more dorsal position of our labeled secondaries while both *alcamb*+ and *bmp16*+ SMns shared a more ventral location (Fig. 6A). We next tested *alcamb+* and *foxb1b+* SMns for potential correspondence to the different subsets of SMns labeled with two transgenic lines, Tg(isl1: GFP) and Tg(gata2: GFP) (Appel et al 1995, Meng et al 1997, Higashijima et al 2000). Labeling with *in situ* probes revealed that expression of *alcamb* overlapped with GFP+ SMns in the Tg(gata2: GFP) fish (Fig. 6B), while *foxb1b* co-localized with those in Tg(isl1: GFP) (Fig. 6C). Thus, *foxb1b*/*alcamb* expression distinguishes previously identified SMn subsets. The third cluster, corresponding to the *bmp* expressing class of SMns, were fewer in number and the targets are not known. However, *bmp* signaling has been linked to specification of slow muscle, raising the possibility that this muscle type receives innervation by *bmp*+ Mns (Kuroda et al, 2013).

**Figure 6.**
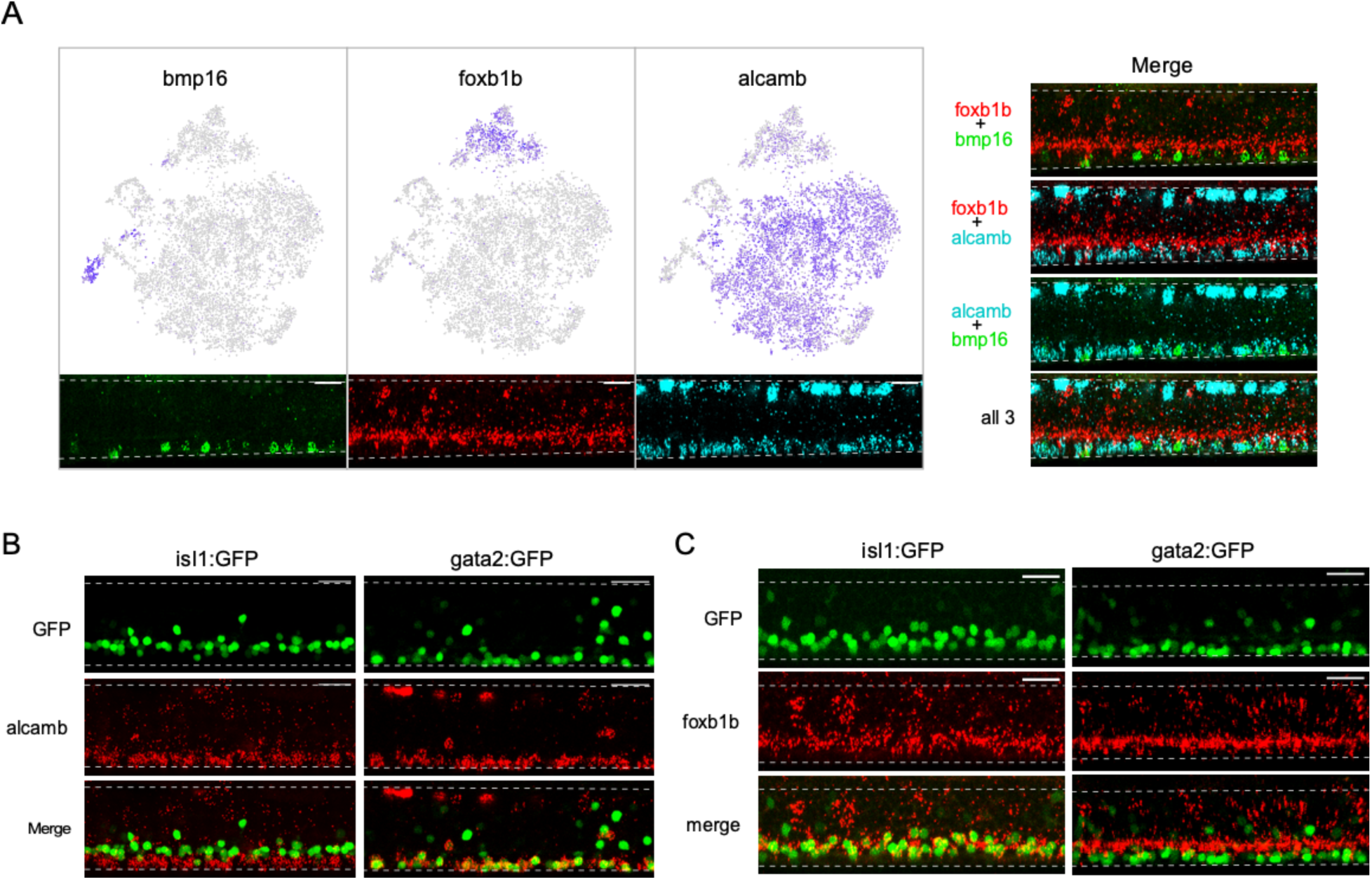
Diversity among SMns. A. Differential expression of *bmp16*, *foxb1b* and *alcamb* associated with distinct SMn clusters. Feature plots in the Mn t-SNE projection (top) and representative *in situ* hybridization images (bottom) are shown for each marker gene. Merged images of different probe combinations (right) highlight the differences in expression pattern in the motor column. n = 10 fish. B. Representative *in situ* hybridization images comparing *alcamb* expression in GFP labeled SMn sub-populations in Tg(isl1:GFP) (left, n = 6 fish) and Tg(gata2:GFP) (right, n = 6 fish). Note that *alcamb* also expresses at high level in the RB neurons located along the dorsal edge of the spinal cord. C. Representative *in situ* hybridization images comparing *foxb1b* expression in GFP labeled SMn sub-populations in Tg(isl:GFP) (left, n=14 fish) and Tg(gata2:GFP) (right, n= 12 fish). Scale bar 20 µm; White dashed line indicates the boundary of spinal cords; Dorsal is up.

### Transcriptome comparisons for the PMn and SMn types

We performed differential expression analysis comparing PMn and SMn transcriptomes in order to identify candidate genes that might account for their functional distinctions previously established using electrophysiology. After applying thresholds to the Mn clusters, based on average levels of gene expression, p-value, and the proportion of cells expressing individual genes, we obtained a list of 508 candidates that showed at least a 30% difference in average expression levels between Mn types, 317 of which were higher in primaries. To guide our identification of genes that play potential roles in governing excitability and synaptic transmission differences between the two groups of Mns (Supplemental Table 2) we used Gene Ontology (GO) enrichment analysis for the PMn specific DEGs (The Gene Ontology Consortium, 2021) (Ashburner et al 2000). We focused on significantly enriched GO terms for DEGs that were associated with 3 broad categories of biological processes, synaptic function, ion channels/transporters and ion homeostasis, and ATP generation (Supplemental Fig. 2), due to their roles in excitability and synaptic transmission. Of the 37 Gene ontology terms identified for the PMn DEGs 24 fell into one of these 3 broad functionally relevant categories (Supplemental Fig. 2). We next manually annotated the molecular function of DEGs by referencing evidence-based database ZFIN (Bradford et al 2022) in order to identify specific functionally relevant genes differentially expressed between PMn and SMn. For the PMn type, DEGs encoding a large number synaptic proteins included those of the core exocytotic machinery such as isoforms of VAMP, SNAP25 and Syntaxin; regulators of exocytosis, including Synaptotagmins, NSF, Complexins and RIM; synaptic vesicle proteins, and synaptic structural proteins such as Synuclein, Bassoon and Piccolo (Fig. 7A). Comparisons of expression levels indicate that no synaptic genes were specifically enriched in SMns, but 29 were enriched in PMns compared to SMns (Fig. 7A). Additionally, 5 synaptic genes were shared at approximately equivalent levels between Mn types (Fig. 7A). Similarly, 8 different voltage dependent ion channel subunits were preferentially expressed in the PMn, with the highest levels corresponding to the P/Q calcium channel, the β4 sodium channel subunit and the Kv3.3 potassium channel. Four other ion channel types were shared by both SMns and PMns. As with the synaptic genes, no ion channel candidates were specifically enriched in the SMn type. The absence of unique ion channel and synaptic gene enrichment in SMns contrasted with the high levels of transcriptional factors and RNA binding proteins, neither of which are likely candidates for conferring functional distinctions between Mn types. Approximately two thirds of the 191 genes enriched in the SMns (118/191) were of these categories. The low levels of candidate ion channels and synaptic genes in the SMns might be expected on the basis of greatly reduced levels of synaptic transmission reflected in quantal content and release probability compared to PMn (Wang & Brehm 2017).

**Figure 7.**
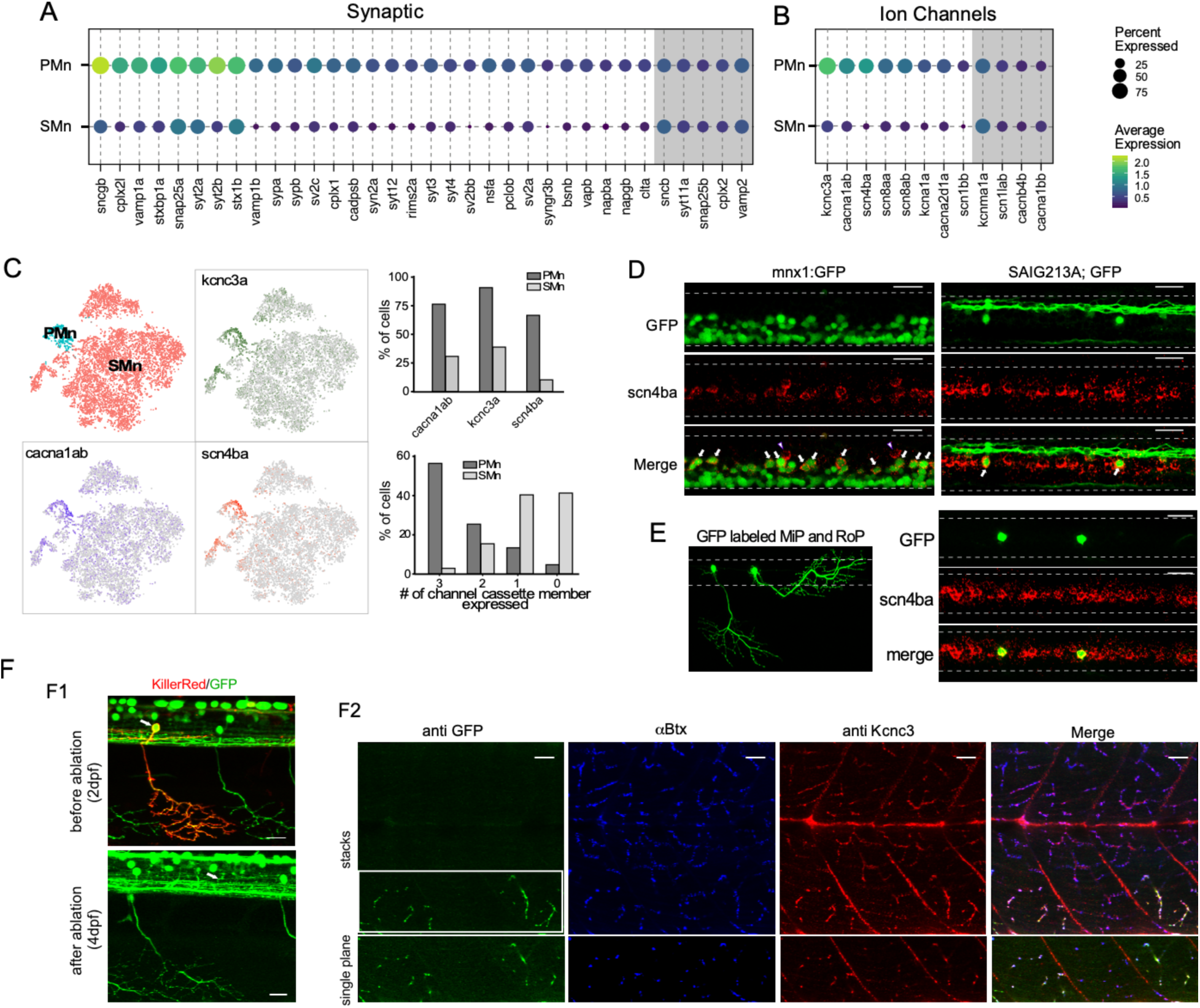
Transcriptome comparison between PMns and SMns. A. Dot plot for synaptic genes differentially enriched in PMns compared to SMns. Both percentage of cells with expression and average expression levels were shown. Examples of synaptic genes that expressed at comparable levels between the two Mn types are shaded gray. B. A similar comparison for differentially expressed ion channel genes as shown in A. C. Feature plots for three top differentially expressed ion channel genes, *cacna1ab*, *scn4ba* and *kcna3a,* shown in the Mn t-SNE projection (left). The assignment of Mn type identity was duplicated from Fig. 5G for reference. (Right graphs) The proportion of cells in each Mn type expressing individual cassette member (top), and cassette member combinations (bottom). D. Representative *in situ* hybridization images with *scn4ba* probes in Tg(mnx1:GFP) (left) and Tg(SAIG213A;GFP) (right) transgenic fish. Each image shows approximately 2 segments of the spinal cord in the middle trunk of 4 dpf fish. Arrows indicate the PMns in Tg(mnx1:GFP) and CaP in Tg(SAIG213A;GFP) fish (n= 15-18 fish). 2 CoLo interneurons labeled with *scn4ba* probes were also indicated (arrowhead). E. Expression of *scn4ba* in the MiP and RoP PMns. The morphology of GFP labeled MiP and RoP in an injected fish shown (left). *In situ* hybridization images with *scnba* probes in this fish showed colocalization with GFP labeling (right). n = 7-10 cells. Scale bar 20 µm; White dashed line indicated the boundary of spinal cord; Dorsal is up. F. Validation of *kcnc3a* enrichment in PMns by immunohistochemistry staining. F1. KillerRed-mediated photo-inactivation of CaP. Representative fluorescent images showing ~ two segments of a Tg(SAIG213A;EGFP) fish with a single CaP (arrow) expressing KillerRed, before photo illumination at 2 dpf (top), and ~40 hrs after inactivation (bottom). Both the soma (the location indicated by an arrow) and periphery branches (see also F2 leftmost panel) are absent after the ablation. F2. Immunohistochemical staining of the same fish with a Kcnc3 specific antibody. GFP expression is revealed by anti-GFP antibody staining, and the location of synapses labeled by α-Btx. Top panels represented a maximal intensity projection of a stacked of z-plane images, while the bottom showed a single focal plane of the CaP target field (indicated by a white box). Scale bar 20 µm. n= 5 fish.

Of particular interest among the genes specific to the PMn type were the trio of highest expressing voltage-dependent ion channel types that have been individually linked in previous studies to augmenting transmitter release and/or AP firing rate. Those channel types include a voltage-dependent P/Q-type calcium channel α subunit (*cacna1ab*), a Kv3.3 potassium channel α subunit (*kcnc3a*), and a sodium channel b4 subunit (*scn4ba*) (Fig. 7B & C) (Eggermann et al 2011, Lewis & Raman 2014, Zhang & Kaczmarek 2016). The identification of the *cacna1ab* channel as a top DEG in PMns corroborated previous studies firmly establishing the P/Q-type as the presynaptic active zone calcium channel mediating Ca^2+^-dependent release specifically in the PMns (Wen et al 2013, Wen et al 2020). Enriched expression of the sodium channel b subunit *scn4ba* in PMns was validated by *in situ hybridization* (Fig. 7D). *scn4ba* probes specifically labeled dorsally-located Mns with large sizes in the Tg(mnx1:GFP) fish (Fig. 7D). These represented the PMns, as shown by co-labeling of individually labeled CaP, MiP and RoP Mns (Fig. 7D & E). The zebrafish *kcnc3a* has been shown to be expressed preferentially in PMn at embryonic ages (Issa et al 2011). We tested its expression at 4 dpf fish using immunohistochemical staining. A *kcnc3* sub-type specific antibody efficiently labeled NMJ synaptic terminal marked by α-bungarotoxin (α-Btx, Fig. 7F2). Since the PMn and SMn axons track along with each other and form synapses with the same postsynaptic receptor clusters (Wen et al 2020), further resolution was needed to distinguish the PMn and SMn synaptic terminals. For this purpose, we ablated individual CaPs by transiently expressing the phototoxic KillerRed protein (Formella et al 2018), followed by light inactivation at 2 dpf. By 4 dpf, the ablation of CaP was complete as indicated by the absence of its soma and processes (Fig. 7F1). *Kcnc3* antibody staining showed a specific signal reduction in synapses located in the ventral-most musculature (Fig. 7F2), the target field shared by the CaP and numerous other SMns. This result strongly supports the expression specificity of *kcnc3a* channel in the PMns.

These specific calcium, potassium and sodium channel isoforms co-express specifically among the four types of PMns. We term this collective unit of three different voltage-dependent ion channel types as a ‘channel cassette’. Over 55% of the cells in PMn cluster express the cassette, compared to <3% in the SMns over all (Fig. 7C). Together with other synaptic DEGs revealed by the analysis, our results suggested a molecular logic for the functional specialization in PMn synapses, such as strong release and high frequency firing that are uniquely associated with escape behavior.

### Gene ensembles for additional components of escape circuitry

Further insights into the potential contribution the ion channel cassette to fast synaptic function specialization came from examination of their differential expression pattern in other circuitry components involved in high-speed swimming and escape. Those cell types include the CoLo inhibitory and v2a excitatory interneurons, both of which provide direct synaptic connections to the PMn type. Both type I and type II v2a neurons participate in high swim speeds, but the type II is more effectively recruited over the range of higher speeds (Hale et al 2001, Higashijima et al 2004c, Kimura et al 2006, Liao & Fetcho 2008, McLean et al 2008, Satou et al 2009, Ampatzis et al 2014, Menelaou & McLean 2019, Satou et al 2020). Additionally, electrophysiological studies have shown that type II v2a neurons fire faster and more reliably than type I v2a neurons (Menelaou & McLean 2019). As seen in the PMn type, the coexpression of the ion channel cassette is high in both type II v2a and CoLo interneuron types compared to other interneuron types (Fig. 8A). Some of those interneuron types expressed individual components of the cassette, most often *kcnc3a* or *cacna1ab* channel types, but not the full cassette. Indeed, the *scn4ba* sodium channel β subunit is the strongest delineator among the 3 for inclusion into the ion cassette classification (Fig. 8A & 7C).

**Figure 8.**
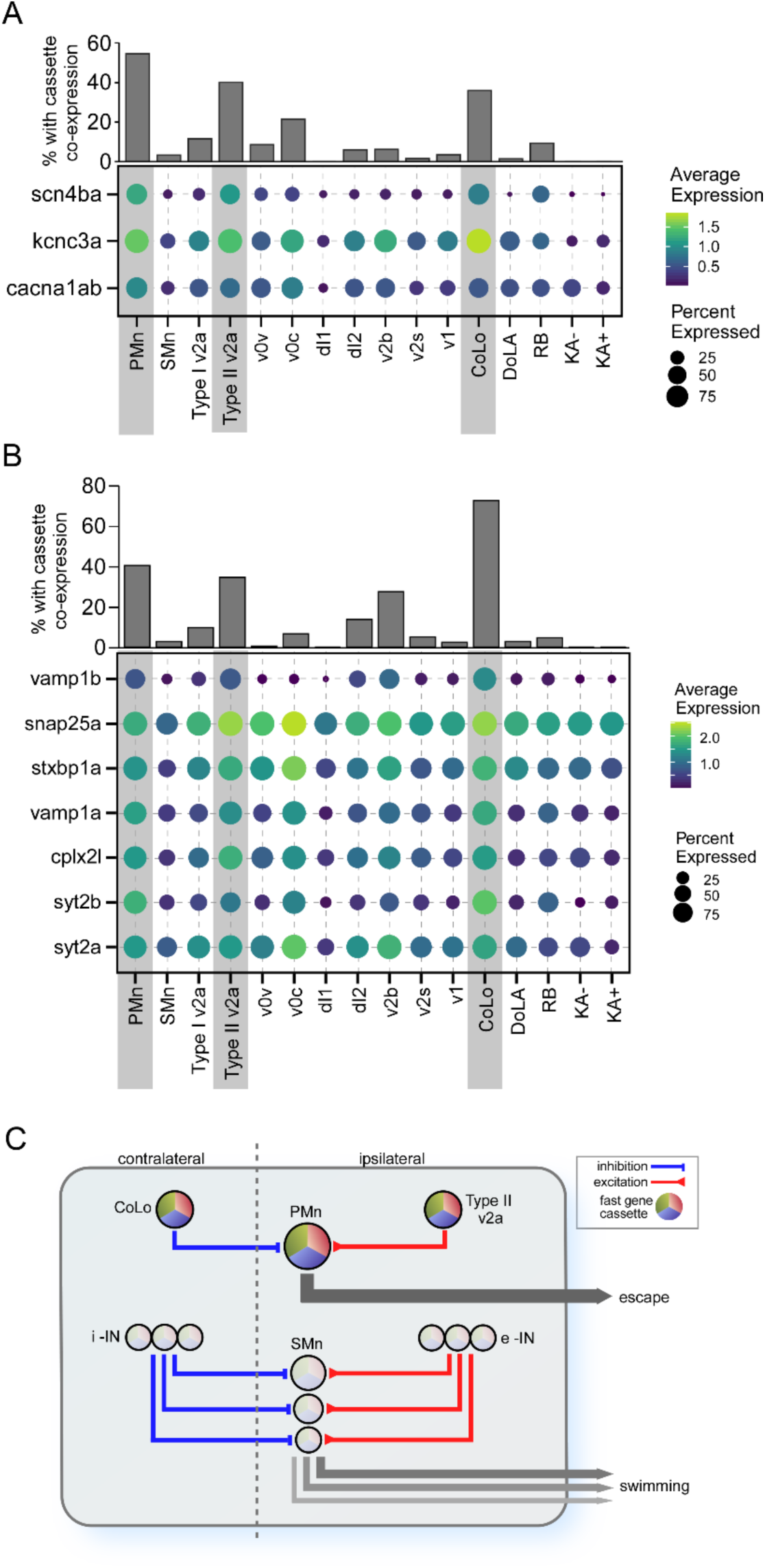
Differential expression of gene cassettes in larval zebrafish escape circuit. A. The ion channel cassette. (lower) Dot plot showing the averaged expression level (color scale) and percentage of cell expressed (dot area) of *scn4ba*, *kcnc3a* and *cacna1ab* channel genes in different neuronal types. The three neuronal types located at the escape circuit output pathway are highlighted (gray shade). (upper) Bar graph showing the percentages of cells in each neuronal type co-expressing all three channel genes. B. The cassette of synaptic genes. Seven top PMn DEGs encoding proteins involved in synaptic transmission were shown for all neuronal type. C. Proposed circuitry for separate control over escape and swimming in larval zebrafish. The schematic model is based on published studies and incorporates the role of the differentially expressed gene cassettes in conferring behavioral and functional distinctions that are manifest both centrally and at the NMJ. The circuitry and cassette expression in the PMn that control escape is illustrated at the top and the SMn circuitry that controls swim speed is illustrated at the bottom. Swim speed is dependent on Mn size as published which is determined at the levels of both spinal circuitry and neuromuscular synaptic strength. According to this simplified model the gradient of synaptic strength and speed at the NMJ is set by the levels of cassette expression among Mns, i-IN: inhibitory interneurons; e-IN: excitatory interneurons.

Top DEGs identified for the PMns that encode synaptic proteins also showed higher levels of expression in Type II v2a and CoLo interneurons compared to other cell types, including *vamp1a, vamp1b, syt2a, syt2b, snap25a, stxbp1a* and *cplx2l*, all central players in transmitter release (Fig. 8B). This further strengthens the proposition that a synaptic cassette works collectively with the ion channel cassette to create a strong neuronal circuitry controlling the escape behavior and fast swimming behavior. These results are incorporated into a simplified model that potentially accounts for the differential circuitry that distinguish regulation of slow swimming from the power circuits (Fig. 8C).

## Discussion

Interest in the spinal circuitry controlling muscle movements among vertebrates remains high, especially from the standpoint of understanding inheritable human myasthenic disorders (Walogorksy 2012, Wen 2016a, Ono 2002, Wang 2008, Downes & Granato 2004). As a model for investigation into spinal circuits, zebrafish offers great simplicity when linking spinal circuitry to a specific behavior. Moreover, morphological, physiological and transgenic labeling techniques, have revealed a large repertoire of spinal neuronal types involved in locomotion in zebrafish, many of them sharing homologies with those in mouse (Grillner 2003, Goulding 2009, Grillner & Jessell 2009). In both mouse and larval zebrafish, the understanding of motility circuitry centers on the study of Mns which are the final pathway to movement. Unlike the numerous Mn subtypes in mammals, which are grouped according to different anatomical positions and muscle targets (Stifani 2014), there are two main classes in zebrafish which share the same fast muscle cells as well as the individual neuromuscular synapses (Eisen et al 1986, Myers et al 1986, Menelaou & McLean 2012, Bello-Rojas et al 2019, Wen et al 2020). To aid in the assignment of neuronal types to specific circuits, we performed scRNAseq analysis on larval zebrafish spinal cord. Our study revealed new markers for key components of the spinal circuity that likely contribute to regulation of different behavioral responses, along with identification of a new interneuron type.

Studies on Mn control of movement in zebrafish have focused on two idiosyncratic swimming behaviors; the single powerful tail bend initiating escape and the subsequent, generally less powerful, rhythmic swimming (Budick & O’Malley 2000, Thorsen et al 2004). The spinal circuits mediating these two behaviors have been the subject of many studies, which conclude that the SMns do not play important roles in behavioral escape but instead regulate swimming over a broad range of speeds. Consistent with this role they cover a lower range of AP firing and release much less transmitter per AP compared to the PMn type (McLean et al 2007, Wang & Brehm 2017, Wen et al 2020). Thus, PMns regulate the highest speed swimming and escape behavior. Indeed, the functional distinctions were our impetus to identify individual transcriptomes for the separate Mn types. Using a newly identified marker, *chaga*, we identified a single small cell cluster corresponding to the four PMns located within each hemisegment. The SMns, by contrast, corresponded to three different clusters based on the markers *foxb1b, alcamb* and *bmp16*. These three transcripts were used to localize SMn subtype populations within the spinal cord. The SMn somas containing *foxb1b* transcripts formed the most dorsal group which are positioned above the *alcamb*-labeled group in the motor column. The smallest cluster, labeled by *bmp16*, overlapped with the *alcamb* label. The relative position of these SMn clusters is consistent with studies showing a correspondence between the dorsal-ventral position within the spinal cord and control of rhythmic swim speed by the SMns. Sequential recruitment of more dorsally located SMns leads to the generation of increased power and faster swim speed (McLean et al 2007, Gabriel et al 2011, Wang & Brehm 2017). This gradient of power is determined through both firing pattern and by amount of transmitter release at the NMJ (Wang & Brehm 2017). Thus, it is plausible that the transcriptomic distinctions among SMns reflect their differential roles in swim speed determination and their functional distinctions in synaptic strength (Wang & Brehm 2017, Wen et al 2020).

Previous electrophysiological studies indicate that PMns fire APs at a high frequency and that transmitter release occurs with a short synaptic delay, high quantal content and a high release probability (Wen et al 2016b). By contrast, SMn synapses are much weaker and variable in firing properties, in keeping with the distinct behavioral roles of PMns and SMns (Wang & Brehm 2017, Wen et al 2020). Consistent with these distinctions, analysis of the transcriptomes for the two Mn types revealed large scale differences. The PMns expressed high levels of transcripts encoding ion channels and exocytotic machinery, all of which are generally not detected in SMns. Overall, the transcriptomic profile comparisons are consistent with the electrophysiological findings of greatly enhanced neuromuscular transmission for the PMns. In particular, the PMns had very high expression of an ion channel cassette formed by three different voltage-dependent ion channel types, each of which had been linked previously to either high release probability of transmitter or very high AP frequency. The cassette transcript with the most restrictive expression pattern encoded the β4 subunit of the voltage-dependent sodium channel NaV1.6. This subunit confers high frequency firing through a fast reversible block of the NaV1.6 channel pore. The blocking kinetics are sufficiently fast to enable the neuron to fire APs at frequencies exceeding the refractory period (Raman & Bean 1997, Grieco et al 2005, Lewis & Raman 2014, Ransdell et al 2017). A second highly enriched voltage-dependent channel in PMns is the Kv3.3 potassium channel. This ion channel type has been associated with high AP firing frequency, as well as with augmented transmitter release in mammalian neurons (Zhang & Kaczmarek 2016, Richardson et al 2022), both due to fast activation kinetics that result in fast repolarization of the AP. The zebrafish *kcnc3a* gene, encoding the Kv3.3 channel, gives rise to a transient potassium current that has both fast activation and inactivation making it well suited for its proposed role in shortening AP waveform and allowing high frequency firing (Mock et al 2010). The third cassette member, *cacna1ab*, encoding a P/Q-type calcium channel, was a top DEG in the PMn, in agreement with our previously published finding that the SMn expresses a different calcium channel isoform, most likely N-type based on sensitivity to specific conotoxin isoforms (Wen et al 2020). Mutations in the *cacna1ab* gene completely abolished AP-evoked release in PMns but left synaptic transmission in SMns intact (Wen et al 2020). As a result, mutant fish are unable to mount a fast escape response, but are still capable of normal fictive swimming (Wen et al 2020). The P/Q type calcium channel has been associated widely with synapses with high release probability (Iwasaki & Takahashi 1998, Wu et al 1999, Stephens et al 2001, Fedchyshyn & Wang 2005, Bucurenciu et al 2010, Eggermann et al 2011, Young & Veeraraghavan 2021), a feature, due in part, to a higher open probability during APs compared to the N-type counterparts (Li et al 2007, Naranjo et al 2015), thereby promoting calcium entry. We hypothesize that the collective actions of this ion channel cassette, with their unique biophysical properties, serve to mediate the escape response and the highest swim speed through ultrafast repetitive firing and maximal release of neurotransmitter. This trio of voltage dependent ion channels is also co-expressed in mammalian neuronal types that are involved in fast signaling, suggesting a conserved role for the cassette in fast behaviors. Examples include the auditory neuron calyx of Held (Iwasaki and Takahashi 1998, Midorikawa et al 2014, Richardson et al 2022, Kim et al 2010) and the pyramidal neurons of the brain (Grieco et al 2005, Akemann and Knopfel 2006, Hillman et al 1991).

A second cassette comprised of a set of genes involved in synaptic function was also revealed by comparing the transcriptomes between the PMns and SMns. The cassette components enriched in PMns were genes encoding isoforms of VAMP, Syntaxin, Synaptotagmin, SNAP25 and Complexin, all components associated with exocytosis and transmitter release. Unlike the ion channel cassette described above, which is strongly linked to synapses with high release probability and/or high firing rate, the actions by which these synaptic genes could differentially support strong and fast synaptic properties remains speculative. It has been suggested, however, at both fly NMJ and mammalian CNS, that the differential synaptic strength among synapses correlates with abundance of proteins involved in transmitter release (Holderith et al 2012, Peled et al 2014, Akbergenova et al 2018).

Further support for the idea that ion channel and synaptic cassettes both play direct roles in formation of specialized circuitry surrounding the PMn was provided by transcriptomic analysis of interneuron types known to interact specifically with the PMn and to be recruited during high speed swimming. Those include the Type II v2a excitatory interneuron and CoLo inhibitory interneuron forming the output pathway for the escape response (Bhatt et al 2007, Liao & Fetcho 2008, Satou et al 2009, Menelaou & McLean 2012, Menelaou & McLean 2019). As with the PMn, both interneuron types coexpress the triple ion channel cassette members at high levels compared to those interneurons that participate less during recruitment during high speed swimming. As shown for distinction among Mn types, the sodium channel β4 appears to be the most restrictive among the three ion channel types in conferring fast firing to the interneurons as well. Unlike the case for the PMn, the AP firing and transmitter release properties of these neuronal types are less well established. However, it is clear that the Type II v2a, in particular, can fire at high frequencies over 600 Hz (Menelaou & McLean 2019). The highly specific enrichment of these two gene cassettes, in neurons involved in escape behavior, suggests that they serve as an integral part of the molecular signature underlying functional specialization. It remains to be seen whether the gene cassettes we identified for zebrafish escape circuit represent a general transcriptional architecture plan to build synapses with great strength and speed in the CNS of higher vertebrates.

Finally, our scRNAseq analyses provides a resource for future identification of gene functions that are causal or associated with human disorders involving Mn dysfunction. In the context of myasthenic disorders in particular, zebrafish has provided a large number of animal models corresponding to human syndromes, including slow channel syndrome, episodic apnea, and rapsyn deficiency (Ono et al 2002, Wang et al 2008, Walogorsky et al 2012a, Walogorsky et al 2012b, Wen et al 2016a). Both myasthenic syndromes and amyotrophic lateral sclerosis (ALS) involve dysfunction at the level of the motor circuits (Ferraiuolo et al 2011). In the case of ALS, it is well known that fast motor neurons are selectively targeted for degeneration (Hadzipasic et al 2014, Nijssen et al 2017). Our analysis comparing transcriptional profiles between fast versus slow motor circuit components offers a new means for probing the transcriptional consequences of neuromuscular disease states.

## Materials and Methods

### Fish lines and husbandry

The transgenic line Tg(mnx1:GFP) was provided by Dr. David McLean (Northwestern University). Tg(vsx2:Kaede) was provided by Dr. Joseph Fetcho (Cornell University). Tg(SAIG213A;EGFP), Tg(islet1:GFP) and Tg(gata2:GFP) were maintained in the in-house facility. Zebrafish husbandry and procedures were carried out according the standards approved by Institutional Animal Care and Use Committee at Oregon Health & Science University (OHSU). Experiments were performed using larva at 4 dpf. Sex of the larva cannot be determined at this age.

### Sparse labeling of spinal neurons

In most cases, we used the Gal4-UAS system to achieve mosaic expression by co-injection of two plasmids into single cell embryos: one containing Gal4 driven by cell type – specific promoters and the other containing fluorescent reporter genes under the control of UAS element. The mnx1 or vAChT promoter drove the expression in Mns (mnx1:Gal4 plasmid provided by Dr. McLean, Northwestern University and vAChT:Gal4 provided by Dr. Joe Fetcho, Cornell University). drmt3a promoter (drmt3a:Gal4 provided by Dr. Shinichi Higashijima, National Institute of Natural Sciences, Japan) was used for expression in glycinergic inhibitory interneurons. mCherry driven by HuC promoter was used to label spinal neurons, including the DoLA interneurons. Injected fish were screened on 3 dpf for sparse fluorescent neurons that could provide detailed morphology.

### KillerRed mediated CaP ablation

We transiently expressed the phototoxic KillerRed protein in Tg(SAIG213A:EGFP) fish by injecting a plasmid expressing KillerRed driven by UAS promoter (Addgene plasmid # 115516; a gift from Marco Morsch). Fish with KillerRed expression in CaP were identified. Individual CaPs were ablated by light inactivation at 2 dpf by illuminating for 10 mins with a 560 nm laser set at high power. This completely bleached KillerRed fluorescence, and induced visible blebbing in CaP terminals. Fish were grown to 4 dpf, and the ablation was confirmed by the absence of GFP-labeled soma and neurites.

### Whole mount immunocytochemistry

Whole mount immunohistochemistry was performed as described previously (Wen et al 2020). 4 dpf larvae were fixed in 4% paraformaldehyde at 4 °C for 4 hrs. Zebrafish Kcnc3a channel was labeled using a polyclonal antibody originally generated against the human Kcnc3 protein (ThermoFisher Scientific PA5-53714) at a concentration of 2.5 μg/ml. Polyclonal anti-GFP (Abcam ab13970) was used at 1 μg/ml. Alexa Fluor-conjugated secondary antibodies (Thermo Fisher Scientific) were used at 1 μg/ml. To mark the location of synapses, 1 μg/ml CF405s-conjugated α-Btx (Biotium) was included in the secondary antibody incubation to label postsynaptic acetylcholine receptors.

### Whole-mount in situ RNA hybridization

Fluorescence *in situ* RNA hybridization (FISH) was performed on whole-mount 4 dpf larva using the multiplexed hybridization chain reaction RNA–FISH bundle (HCR RNA-FISH) according to the manufacturer’s instructions (Molecular Instruments) (Choi et al 2016). Probe sets for zebrafish *alcamb, nr2f1a, chga, scn4ba, foxb1b, bmp16, gjd2b* and *pnoca* were custom-designed based on sequences (Molecular Instruments) and used at 4 nM each. Fluorescent HCR hairpin amplifiers were used at 60 nM each to detect the probes. GFP and mCherry fluorescence in the transgenic lines and transient labeled neurons survived the FISH protocol with signal loss mostly limited to the periphery. Residual fluorescence was sufficient to mark the location of soma in the spinal cord without the need for additional amplification.

### Fluorescence imaging

After staining, fixed larval were mounted in 1.5% low melting agarose and imaged on a Zeiss 710 laser-scanning microscope equipped with an LD C-Apochromat 40x/1.2 n.a. objective. Z-stacks of confocal images were acquired using Zen (Carl Zeiss) imaging software, and presented as either maximal intensity projection or single focal planes as indicated in the figures (ImageJ, National Institutes of Health).

### Single cell suspension from spinal cord for scRNAseq

Single cell suspensions were prepared from Tg(SAIG213A;EGFP) fish for the two full spinal cord datasets, and from Tg(mnx1:GFP) for the two FACS-sorted Mn enrichment datasets. About 150 4 dpf larva were euthanized in 0.02% tricaine, and individually decapitated behind the hindbrain. They were incubated with 20 mg/ml collagenase (Life Sciences) in a buffer containing 134 mM NaCl, 2.9 mM KCl, 1.2 mM MgCl_2_, 2.1 mM CaCl_2_, and 10 mM Na-HEPES (pH 7.8) at 28 °C for 2 hr, with intermittent trituration using a p200 pipette aid at 0, 0.5 hr and 1 hr of the incubation. To release spinal cords from remaining tissue, the final triturations were done using fire-polished Pasteur pipettes with decreased opening sizes (300, 200, 100 μm respectively). Intact spinal cords were transferred to L15 media and washed 3 times with fresh media. The spinal cords were incubated with 0.25% trypsin solution (in 1xPBS containing 1 mM EDTA) at 28 °C for 25 min. The digestion was terminated by adding 500 μl stop solution (L15 with 1% fetal bovine serum). The tissue was collected by spinning at 400 g for 3 min at 4°C, washed once with L15 and resuspended in 200 μl of L15 media. Spinal cord cells were dissociated by triturating the digested tissue with fire-polished Pasteur pipettes with 80-100 μm opening. The solution was filtered through a 35 μm strainer into a siliconized collection tube. The suspension was examined on a microscope for cell count, Trypan blue staining based - viability test and proportion of dispersed single cells. Samples with a viability above 70% were used for sequencing.

### FACS sorting

Single cell suspension prepared from Tg(mnx1:GFP) fish were FAC sorted for EGFP+ cells using a 100 μm nozzle on a BD inFlux cell sorter (Flow Cytometry Shared Resource, OHSU). Cells were collected in 100 μl PBS containing 0.2% bovine serum albumin in a siliconized tube.

### Single cell capture, cDNA synthesis, library preparation and sequencing

Single cell capture, cDNA synthesis and library preparation were performed by the Massive Parallel Sequencing Shared Resource at OHSU using the 10x Genomics Chromium v3.0 reagent kit. Single cell suspension for the two full spinal cord replicates targeted 10,000-15,000 cells. For the two FACS sorted samples 4,000-5,000 cells were targeted. Replicate samples were prepared from different clutches of animals. Libraries were sequenced on an Illumina NovaSeq 500 instrument to an average read depth of ~40,000 per cell.

### Reference genome generation and alignment

Cellranger v6.1.1 (10X genomics) was used for the reference genome generation and alignment. Our reference genome was generated by modifying a preexisting reference genome, Lawson v4.3.2 (Lawson et al 2020), which we edited to add an EGFP sequence as an artificial chromosome and to correct a selection of gene names and 3’ untranslated region (UTR) annotations. A full account of all changes made to the Lawson reference genome can be found in Supplemental Table 3, with a representative example shown (Supplemental Fig. 3). Alignment to this reference genome was performed using Cellrangers count function with expect-cells set to the targeted number of cells for each replicate.

### Preprocessing, normalization, clustering analysis and visualization

Count matrices were processed using the Seurat v4.0 package for R (www.satijalab.org/seurat/) (Hao et al 2021). Genes with expression in less than three cells were excluded from further analysis. Initial quality control was performed on each sample independently. Cells were kept for further analysis if they had a number of unique genes between 400 and 4000, UMI counts between 1500 and 9000, and <5% mitochondrial gene content. No further doublet removal methods were applied. Data was normalized using the Seurat Sctransform v2 package in R, generally following the procedure outlined in the Introduction to SCTransform v2 regularization vignette (Hafemeister & Satija 2019, Choudhary & Satija 2022). Principal components were calculated, followed by nearest neighbor graph calculation using ANNoy implemented through Seurat. Clustering used the Leiden community detection method implemented with the FindClusters function and the Leidenalg Python package (Traag et al 2019). To visualize clustered data sets we used t-distributed stochastic neighbor embedding (t-SNE) (van der Maaten & Hinton 2008) or uniform manifold approximation projection (UMAP) (McInnes & Healy 2018) implemented through Seurat.

### Dataset integration

Datasets were combined using Seurat’s integration pipeline (Stuart et al 2019), considering 9,000 variable genes. To examine the correspondence between duplicates, exploratory clustering was done in the combined datasets (Supplemental Fig. 4A & B). Intermixing of data points from different samples was inspected both visually, and by plotting the distribution against one another (Supplemental Fig. 4A & B). For the two full spinal duplicates, one cluster of cells exhibited a strong bias towards a single replicate, with over 90% of the cells sourcing from a single replicate (Supplemental Fig. 4A). These cells were not included in the combined dataset for downstream analysis to control for technical and biological variability. The FACS sorted duplicates had no obvious outliners (Supplemental Fig. 4B).

The combined motor neuron dataset was generated by integrating Mns extracted from the full spinal dataset and the FACS sorted dataset, based on expression of canonical Mn markers (Fig. 5A & B). While the relative populations of SMns did vary between the two sample sources, there was no Mn subtypes that could not be identified independently in both methods of Mn isolation (Supplemental Fig. 5A). Differential expression analysis performed independently with either Mn source yielded highly reproducible sets of the DEGs (Supplemental Fig. 5B), further validating the results from the combined dataset.

### Annotation of cell clusters

Specific cell types in the neuronal subset of the spinal data were annotated on the basis of significantly differentially expressed genes. Exploratory clustering analysis was first conducted to filter out cells not of spinal cord origin. One small cluster in the whole spine dataset (0.6% of cells), expresses *tph2/ucn3l/slc18a2* at high level compared to all the rest of the clusters (Supplemental Fig. 6), marker genes for serotonergic raphe nucleus in the hindbrain (Oikonomou et al 2019, Bradford et al 2022). They reflected a trace amount of hind brain tissue during spinal cord dissection, and were removed from the analysis. Contaminating muscle cells were also removed based on expression of *my1pfa/tnnt3b/actc1b* myosin/troponin/actin genes (Supplemental Fig. 6). Overall, these contaminants accounted for ~2.1% of the total cell population isolated from the spinal cord. No cells in the FACS sorted data set were identified as originating from contamination from outside the spinal cord.

### Data analysis and statistical tests

Differential gene expression was calculated using a Wilcoxon rank sum test implemented with the FindMarkers or FindAllMarkers functions in Seurat. Genes were considered to be enriched in a cluster if they had a log2 fold change > 0.38 (corresponding to > 30% enrichment in expression level), expression in at least 30% of cells in one of the groups being compared, and a Bonferroni adjusted p value of < 10^-5^. A more conservative adjusted p value of < 10^-10^ was used for comparison between PMns and SMns.

### Gene ontology analysis

DEGs comparing PMns and SMns were used to generate lists of gene ontology (GO) terms enriched in the PMns using the EnrichR package in the aspect of “biological processes” (Chen et al 2013, Kuleshov et al 2016, Xie et al 2021). GO terms were considered significantly enriched with adjusted p value <0.05.

### Data Accessibility

Raw fastq files and unprocessed aligned data can be accessed for free through the Gene Expression Omnibus (accession number GSE232801). All code used in data processing and figure creation has been deposited, and is freely available on github (https://github.com/JimmyKelly-bio/Single-cell-RNA-seq-analysis-of-spinal-locomotor-circuitry-in-larval-zebrafish).

## Acknowledgments

The authors thank Drs. Apiar Saunders and Alex Nechiporuk for advice with the scRNAseq analysis, Massive Parallel Sequencing Shared Resource at OHSU, for single cell capture, cDNA synthesis and library preparation and sequencing, and Flow Cytometry Shared Resource at OHSU for FACS sorting. Kara Grist provided expert zebrafish husbandry. This research was funded through a grant from the NIH (NS105664) to P.B.

---

Supplemental Table 1. DEGs in spinal neuron clusters.

Supplemental Table 2. DEGs in PMns versus SMns.

Supplemental Table 3. Changes to the Lawson v4.3.2 reference genome.

