## Supplemental Table 3 for "Single cell RNA-seq analysis of spinal locomotor circuitry in larval zebrafish"

Supplemental Table 3. Changes to the Lawson v4.3.2 reference		
gene name	issue	change
adgra1b	"Not present, two copies of adgra1a"	renamed adgra1a on chr13 51054810-51125683 to adgra1b
artna	last exon is included as itÕs own gene	duplicative gene from chr6 3459026-3459340 removed
brt16	single gene split into two genes with the same name 	two genes one from chr 15 18189324-18200358 and the other from chr15 18198878-18209723 are replaced with one gene from chr15 18189324-18209723
btr18	transcript at chr19 3747061-3759142 is a duplicate of btr22 	New gene runs from chr19 3784726-3790595
btr24	extended across two transcripts 	New gene runs from chr19 4051012-4057870
CABZ01066312.1	complete replicate of si:cabz01066312.1	removed from annotation
CABZ01043953.1	two copies of the gene are present. One is carried over from ensemble and overlaps with a totel of three genes 	"gene idÊLL0000004118 removed from annotation 
"
LOC10817961	was one of two genes called sinhcafl 	renamed to LOC10817961 from sinhcafl. location at chr7 29053217-29062157
COX3	name doesnÕt follow mitochondrial naming scheme 	renamed to mt-co3 at chrM 9735-10520
atp2b4	not included in the annotation due to a misnaming error	renamed from atp2b2 to atp2b4 at chr11 22920587-23080970
cplx1	improperly named cplx2	renamed from cplx2 at chr14 51826150-51856895
XM_021479216	improperly named slc37a4b	renamed to XM_021479216 at chr 10 39133536-39134599
XM_021479503.1	improperly named slc37a4b	"renamed to XM_021479503.1 at chr 10 9276052	9285174"
scaf4a	3ÕUTR region runs too long and overlaps almost completely with the sod1 gene	scaf4a nor runs from chr10 25700711-25716953
syt7a	3Õ UTR region misses a large selection of reads including a pollyA tail 	syt7a extended to chr25 4210280-4313699
syt6a	3ÕUTR region reduced due to the improperly extended 3ÕUTR of olfml3a. 	changed the 3ÕUTR to match the refseq annotation syt6a now runs from chr23 24799876-24856025
syt2b	not named in the annotation. LOC559637 matches ncbi deposit for syt2b and deposited sequence matches too.	renamed from LOC559637 to syt2b
syt2b	"3ÕUTR too short, both pollyA and  junctions clearly indicate that the set of 3Õ reads goes further"	"extended 3ÕUTR now sty2b runs from chr6 55101448-55208884	"
manbal	"Transcript improperly extended on the 5Õ end, and 3Õ end"	reference reduced to the transcripts in refseq and ensemble. runs from chr6 55212113-55218906 
XLOC_036173	placeholder name for a segment of reads that are now assigned to syt2b	removed from annotation
BX005254.2	complete duplicate of hox10ca	removed form annotation
dlg4a	first three exons truncated 	exons added back in gene now runs from chr5 68527850-68755158
dlg4b	3Õutr misses the polly A region 	extended 3ÓUTR gene now runs chr23 44263986-44335750
cacna1ba	3ÕUTR misses pollyA region 	extended 3ÕUTR gene runs from chr5 29293580-29488881
cacna1bb	3ÕUTR misses pollyA region 	extended 3ÕUTR gene runs from chr21 14376858-14631395
cacna1ea	3ÕUTR misses pollyA region 	"3ÕUTR extended gene now runs from 14710542-14831580	"
BX544876.2	transcript has only two exons that are duplicates of exons in cacna1sb 	removed from annotation
cacna2d2a	"3ÕUTR misses polly A region, XLOC-036961 placed as a portion of it and has large degree of overlap with cacna2d2a "	"3ÕUTR is extended, gene now runs from  53438500-53911301"
XLOC-036961	"placed inside the 3ÕUTR for cacna2d2a, has "	removed from annotation
cacnb3a	lawson extends the 3ÕUTR but not far enough 	gene runs from chr23 14097478-14191282
cacnb3b	lawson extends the 3'UTR but incorrectly assigns it as a transcript of the neighboring gene 	extended 3ÕUTR gene runs from chr23 26936372-26967320
CU862020.1	transcript overlaps with cacnb3b and md1b with no basis in the reads. only present in ensemble and lawson but not refseq	removed from annotation
scn1lab	3'UTR misses polly A region and junctions links to unmapped 3ÕUTR	3ÕUTR extended gene runs from chr6 10245080-10321670
scn5lab	3'UTR misses polly A region 	gene runs from chr 24 40905100-41153125
kcnd1	missing a chunk of 3Õ UTR that is present in refseq	gene runs from chr8 9866351-9988543
kcnh2b	name is incorrectly set to CABZ01076968.1 	renamed to kcnh2b
kcnj1b	short region of 3ÕUTR lies outside the annotation	"gene runs from chr 10 39294801	39308020	"
si:ch211-286c4.6	incorrectly named as kcnj11l	renamed to si:ch211-286c4.6
kcnj11	lawson has two expanded transcripts that go in opposite directions 	"changed to match refseq and ensembl, gene runs from chr 25 22853718-22854863 "
si:dkeyp-93a5.2	improperly extended to two locations 	"gene now runs from chr25 18722796-18739924	"
XM_009298067.3	improperly assigned as a transcript of si:dkeyp-93a5.2	renamed to XM_009298067.3 to match refseq gene runs from chr25 25092874-25095771
si:dkeyp-67e1.6	improperly extended to two locations 	gene now runs from chr25 15494040-15496485
XM_021470438.1	improperly assigned as a transcript of si:dkeyp-67e1.6	"renamed to XM_021470438.1 to match refseq gene runs from chr25 28583239-28587896	"
kcnn1b	3'UTR misses polly A region 	gene runs chr8 13239582-13336302
GFP	Added EGFP as an extra chromosome 	location GFP 1-900
