## Supplemental Figures for "Single cell RNA-seq analysis of spinal locomotor circuitry in larval zebrafish"

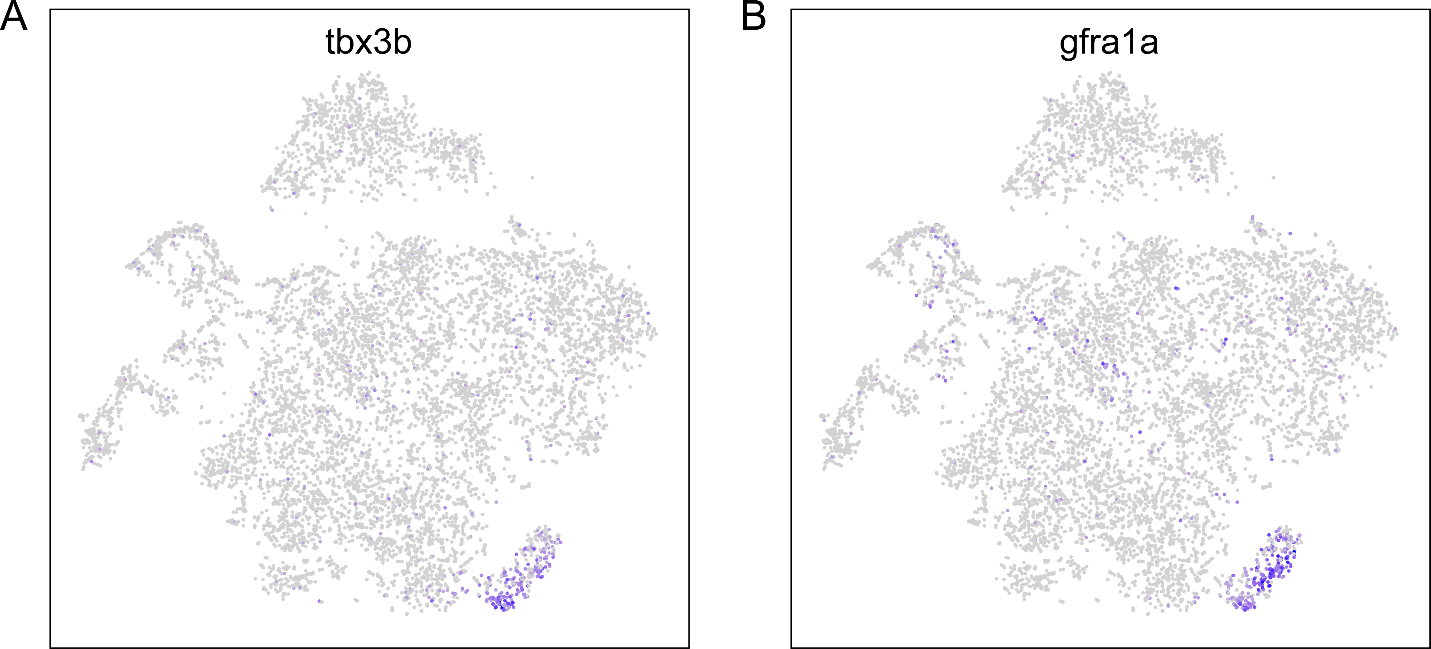

Supplemental Figure 1. An unidentified cluster in the integrated Mn datasets. Feature plots showing restricted expression of *tbx3b* (A), and *gfra1a* (B) to a small cluster in the t-SNE projection of Mn dataset as shown in Fig. 5C.

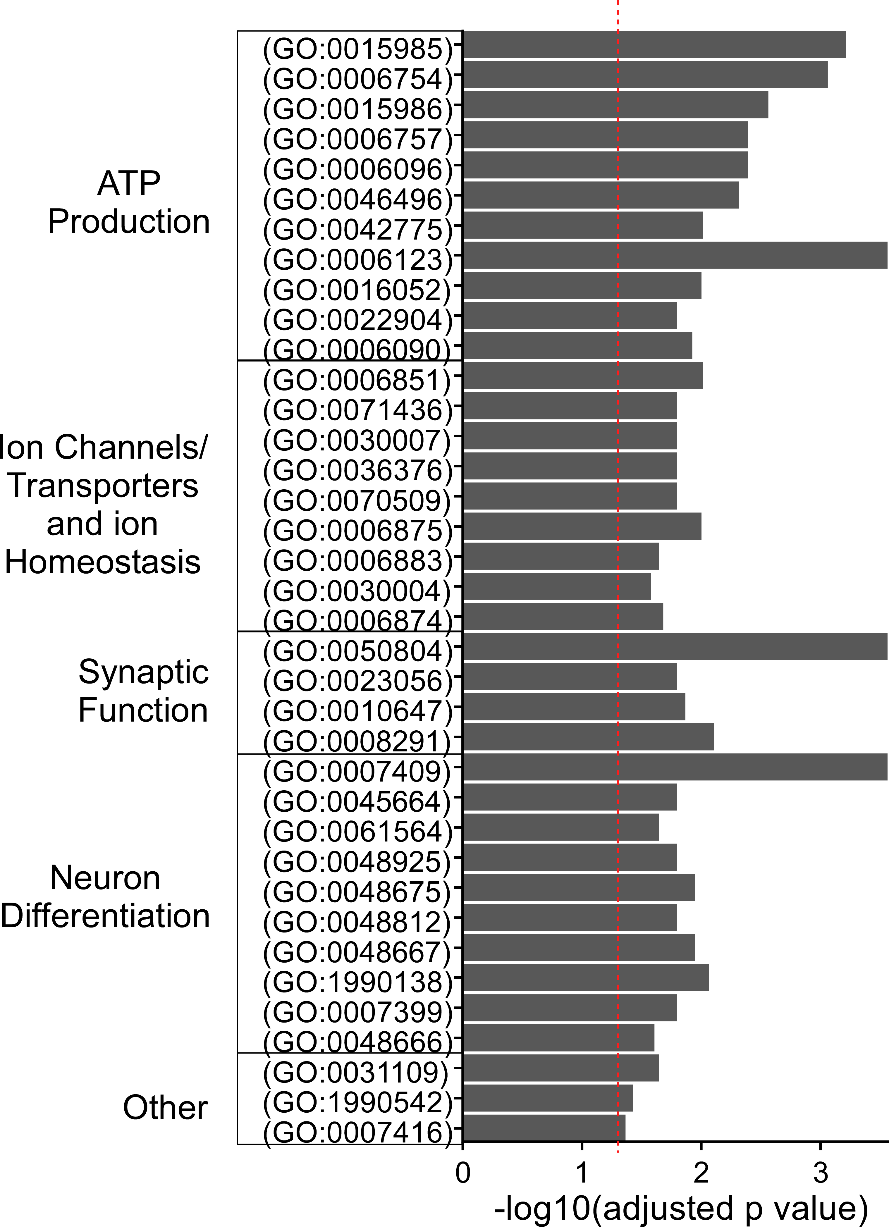

Supplemental Figure 2. Gene Ontology analysis for PMn DEGs. GO terms with significant enrichment for PMn DEGs are shown (red dashed line indicates adjusted p value of 0.05). GO terms are grouped into broad categories, ATP production (11/37), Ion channel/transporters and ion homeostasis (9/37), Synaptic function (4/37), Neuron differentiation (10/37), and other miscellaneous terms (3/37).

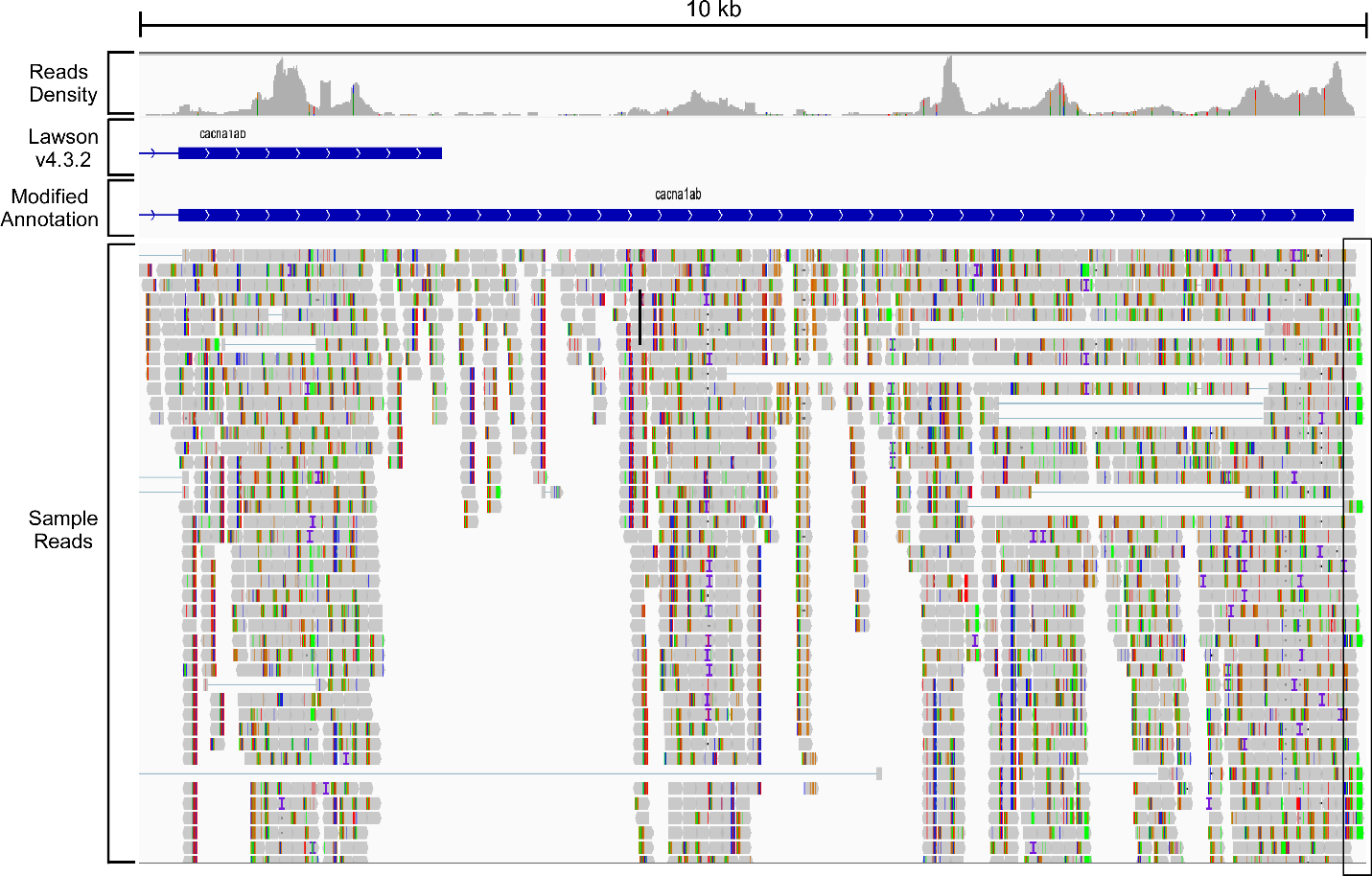

Supplemental Figure 3. Improved annotation of the *cacna1ab* gene to extend 3’ UTR. Select IGV fields shown for 10 Kb of the 3’ region of *cacna1ab,* including the reads density, the annotation based on Lawson v4.3.2. reference genome, the annotation based on our modified reference genome, and sample sequencing reads. Our assignment for the end of 3’ UTR (boxed region) was based on the position of polyA tail in the reads (nucleotide A shown in green).

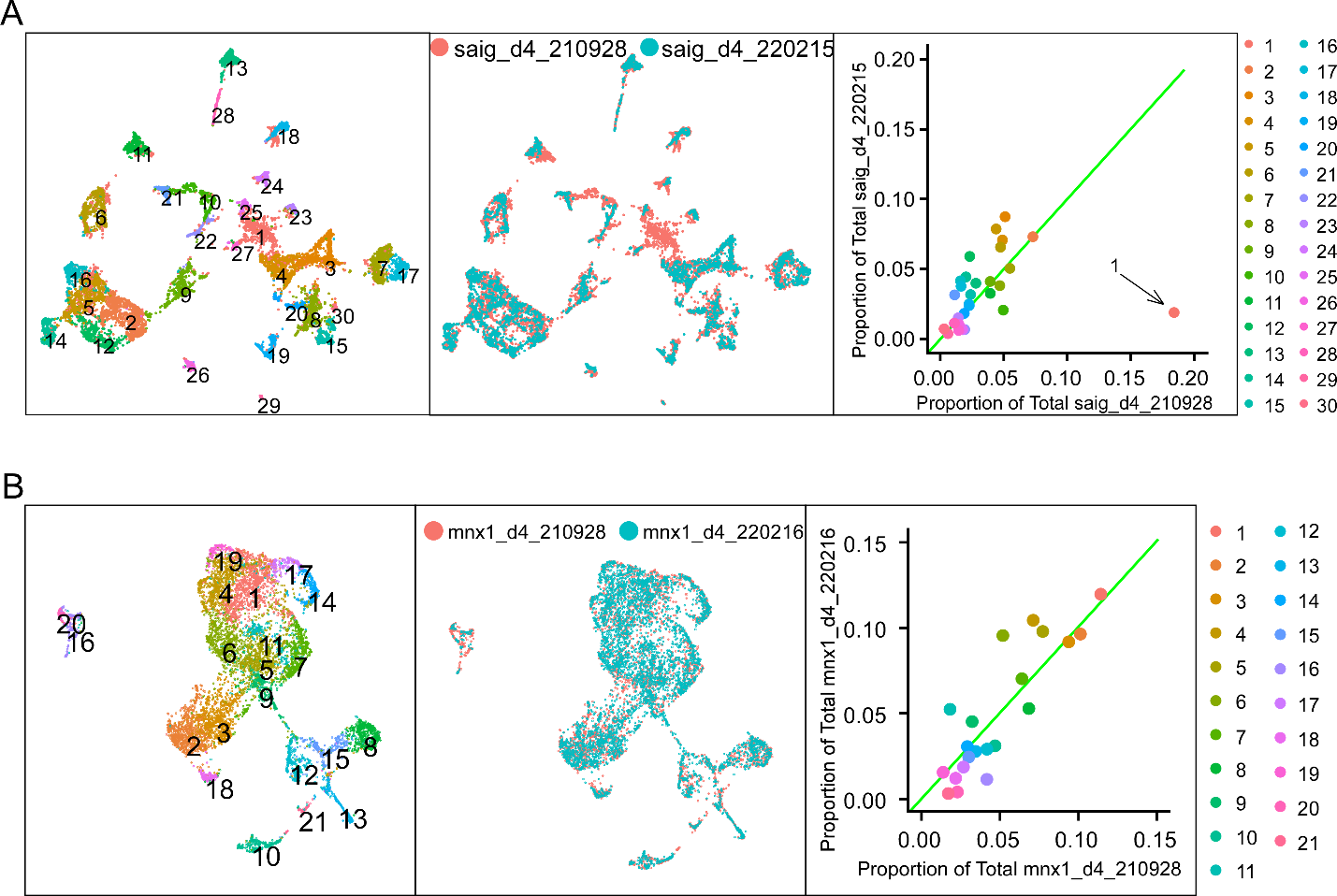

Supplemental Figure 4: Integration of datasets from duplicated experiments. A. Integration of the two full spinal datasets. UMAP plots showing the initial clustering of the integrated full spinal data (left) and the projection colored by sample source (middle). The scatterplot shows the proportion of cell of each source accounted for in each cluster (right). Green line indicates where clusters containing an equal proportion of each source would fall. Cluster 1 is a clear outlier with over 90% of its cells belonging to the saig_d4_210928 sample, and was not included in the final integrated dataset. B. Integration of the two FACS sorted Mn datasets. Correspondence between duplicates was shown similar to A.

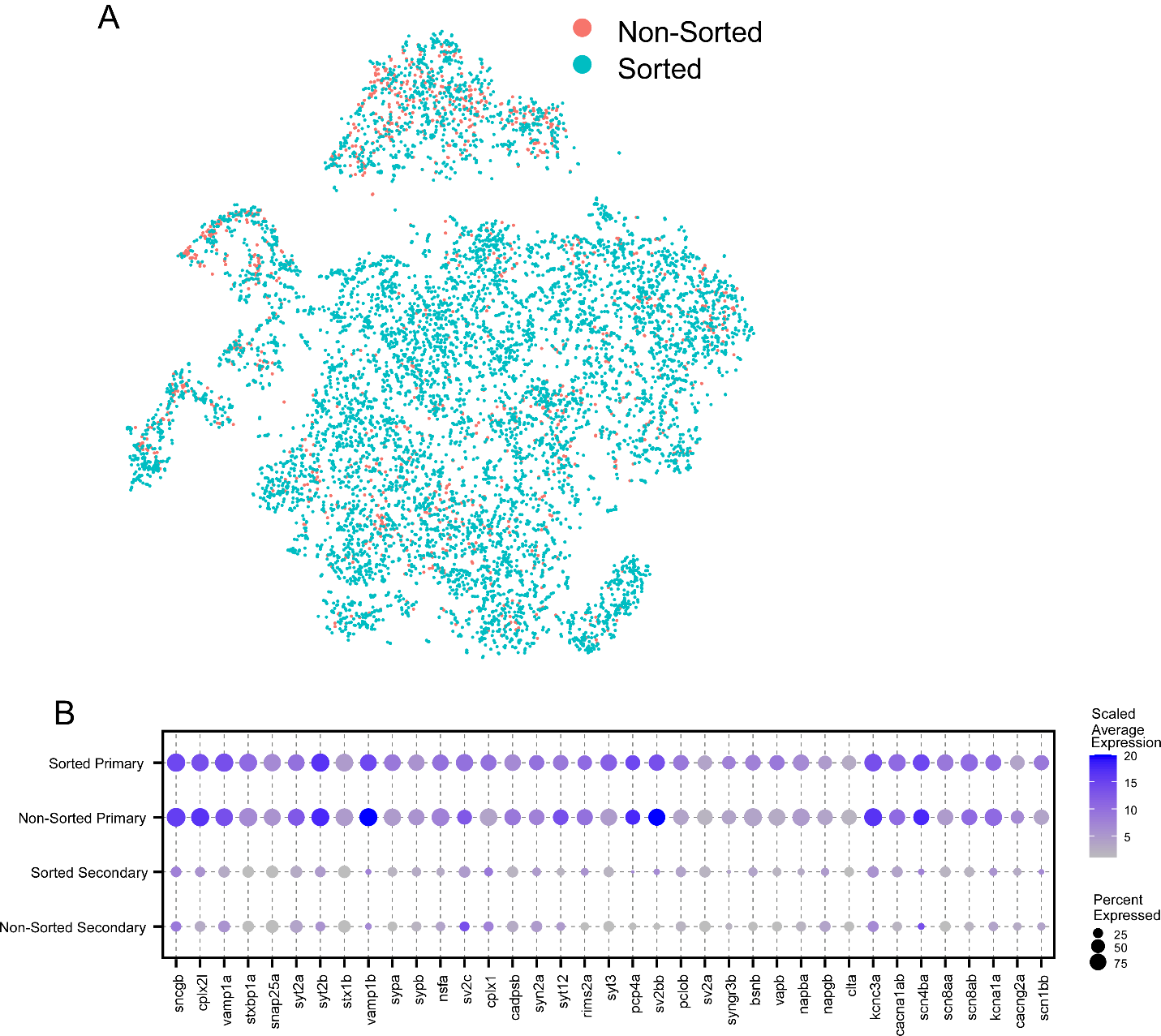

Supplemental Figure 5: Combining Mn datasets from two isolation methods. A. t-SNE projection of the integrated Mn dataset (as shown in Fig. 5C) with cells colored by whether they were sourced from the FACS sorted dataset (sorted, in teal) or extracted from the full spinal dataset (non-sorted, in orange). B. Dotplot comparing results of differential expression analysis between PMns and SMns, done separately for the sorted dataset or the non-sorted dataset. Average expression level was scaled to a range of 1-20.

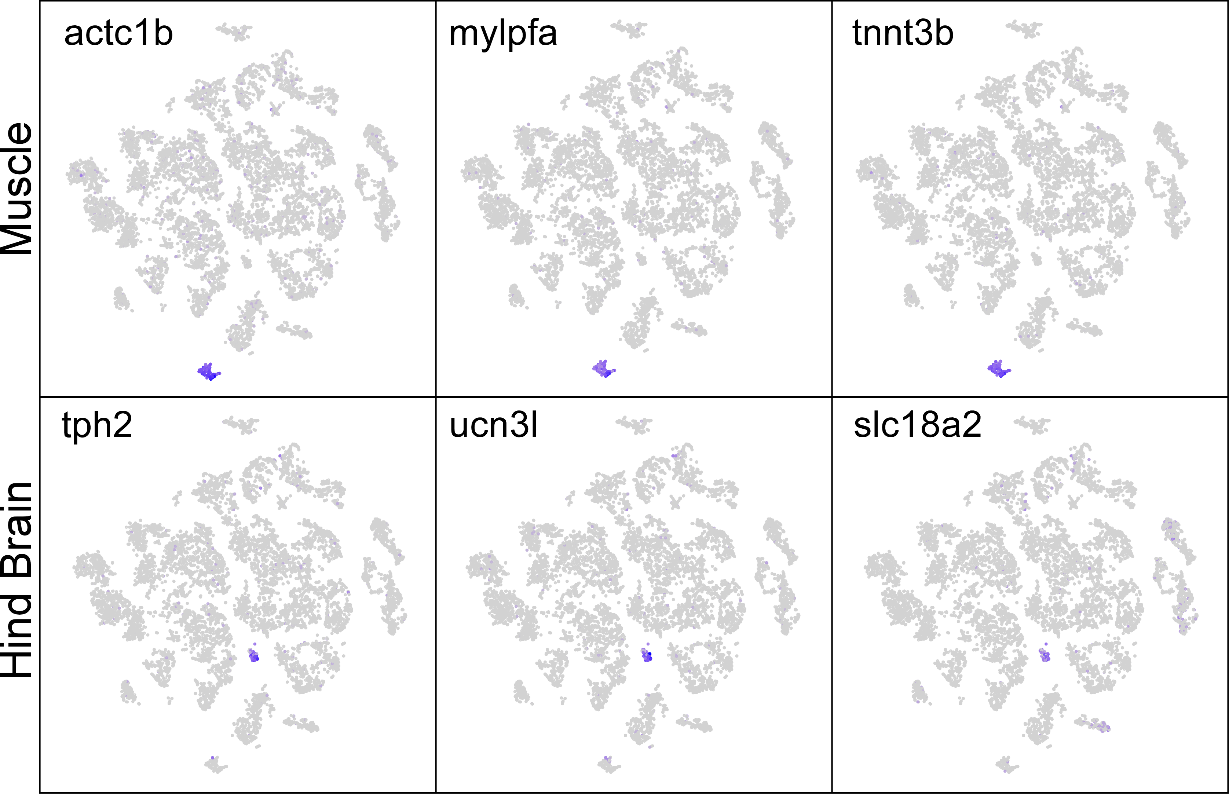

Supplemental Figure 6. Removal of non-spinal cells. Feature plots of the muscle markers (*actc1b, mylpfa, tnnt3b*) and hindbrain markers (*tph2, ucn3l, slc18a2*) in a t-SNE projection of the combined full spinal cord dataset after quality filters. These small clusters were removed on the basis of their non-spinal origin, and not included in the analysis.
